# Phylo-taxonogenomics of 182 strains of genus *Leuconostoc* elucidates its robust taxonomy and biotechnological importance

**DOI:** 10.1101/2021.07.20.453025

**Authors:** Sanjeet Kumar, Kanika Bansal, Santosh Kumar Sethi, Birendra Kumar Bindhani

## Abstract

Genus *Leuconostoc* is a group of a diverse range of lactic acid bacteria (LAB) majorly found in dairy, food and environmental ecology. These microbes are commercially important for several industries due to their intrinsic genomic attributes such as bacteriocins, carbohydrate-active enzymes, plasmids etc. Even though the species of *Leuconostoc* are commercially significant, their taxonomy is largely based on old, low resolution traditional methods. There have been several taxonomic reclassifications in the past which are inadequate for microbiologist and food industry professionals to truly demarcate any new strain of genus *Leuconostoc*. The current taxonomy of the genus is largely based on classical approaches, which are in utmost need of reinvestigation by whole genome-based approaches. In the present study, the taxono-phylogenomic analysis clearly depicted sixteen species including three novel genomospecies in addition to several reshufflings across the species namely, *L. mesenteroides, L. pseudomesenteroides, L. gelidum* and *L. lactis*. The presence of a wide range of carbohydrate-active enzymes, type III polyketide synthase and vector plasmids suggested the biotechnological potential of constituent strains of the genera. Further, the absence of antibiotic gene clusters reaffirms their utility in industries such as food and dairy. Such large-scale in-depth genome-based study can shed light on the nature of the genome dynamics of the species and help to obtain a more robust taxonomic classification.

## Introduction

The genus *Leuconostoc* [1] belongs to the family *Leuconostocaceae* [2] which is one among the widely used group of lactic acid bacteria (LAB). *Leuconostoc* was first isolated by Cienkowski from a slime outbreak in a sugar factory in 1878 [3]. The isolate was named *Ascococcos mesenteroides* which produces dextran slime in sucrose solution [3, 4]. In 1911, an aroma bacteria “X” was isolated from a creamery starter which was later named *Leuconostoc* in 1930 [3, 5]. Historically, the taxonomy of *Leuconostoc* is a long tug of war across the several families of LAB. *Leuconostoc* sp. often occurs in similar habitats as *Lactobacillus* and *Lactococcus*, and was considered as an intermediate between *Streptococcus* and *Lactobacillus*. Genome based investigation of *Lactobacillaceae* and *Leuconostocaceae* suggests union of these two families [6]. In accordance with the latest standing in nomenclature of *Leuconostoc* (https://lpsn.dsmz.de/genus/leuconostoc), there are 16 valid species and along with 11 synonyms. Species of genus *Leuconostoc* are facultatively anaerobic, Gram-positive, nonmotile, catalase-negative, asporogenous, psychrotolerant or psychrotrophic bacteria with optimum growth temperature of 25-30 °C with an average GC content of 37.5% [2]. These are primarily associated with plant matter, fermenting vegetables, industries such as dairy, food and pharma etc. [4, 7–10]. Few reports of their presence in chilled stored meats and human blood are a matter of concern [11, 12]. *Leuconostoc* have been known as a component of starter cultures in dairy since the 1920s, but its factual information about dairy is not very well documented [13]. These microbes are especially, known for their probiotic properties and ability to catalyse the production of several biotechnologically important products [14, 15]. *L. mesenteroides* is the type species of genus *Leuconostoc* [1] which is explored quite extensively in industries. *L. mesenteroides* species are used as a probiotic candidates which enables them to survive and grow under various stress conditions present in the gastrointestinal tract [16]. Species of Leuconostoc is an ideal fermenter of simple carbohydrates and have the capacity to metabolize a wide range of carbohydrates, sugar alcohols and gluconate [17, 18]. Although the *Leuconostoc* sp. has been identified for safe use in the food industry and accorded “generally recognized as safe” (GRAS) status, there are several reports of its disease-causing nature [19, 20]. Interestingly, the strains of *L. mesenteroides* follow dual lifestyles [16] causing disease in plants and humans [21–24].

Even though the species of the genus *Leuconostoc* are widely used for several purposes, the taxonomy is largely unexplored. All the earlier attempts to improve the taxonomy of genus *Leuconostoc* were based on limited phenotypic based methods or lower resolution phylogenetic approaches including housekeeping genes such as16S rRNA, *rpoB, recA* gene etc. [25–28]. As these are based on very limited information, such methods do not provide a robust classification [29]. These studies largely lack a whole genome-based approach to obtain a robust taxonomic, identification of the key functional attributes across the several species of genus *Leuconostoc*. Although, there are some genome based study such as species of *Leuconostoc gasicomitatum* [30] was emended to *Leuconostoc gelidum* subsp. *gasicomitatum* [31]. Later *Leuconostoc mesenteroides* subsp. *suionicum* [32] was emended to *Leuconostoc suionicum* [33]. Most recently genome-based identification of *L. falkenbergense* sp. nov. [34] using whole genome-based sequence information from previously reported strains of *L. pseudomesenteroides*. Another recent study suggests the reclassification of *L. mesenteroides* MTCC 10508 as a strain of *L. suionicum* [35]. All the studies Are focusing on limited strains from some species of Leuconostoc and not considering the whole genus Leuconostoc.

To obtain robust taxonomy of *Leuconostoc* sp., we reinvestigated and re-evaluated the phylogeny, genetic relatedness, and genomic determinants of the species within the present genus *Leuconostoc* (https://lpsn.dsmz.de/genus/leuconostoc). Implementation of these methods resulted in a robust taxonomic framework. We have identified three novel genomospecies namely, GS1 GS2 and GS3. We observed several large reclassifications across the strains of species *L. gelidum* subsp. *gasicomitatum*, *L. pseudomesenteroides, L. lactis* and *L. mesenteroides*. We have also identified species-specific unique genes in the pangenome of the genus *Leuconostoc*. We identified the presence of type III polyketide synthase system in all representative strains *Leuconostoc*, which signifies the antimicrobial ability against pathogenic bacteria. The abundance of the carbohydrate-active enzyme was also identified which includes majorly glycoside hydrolase and glycosyltransferase responsible for degradation, modification, and creation of glycosidic bonds. Furthermore, we believe such a large whole genome-based investigation can help in the estimation and evaluation of new potential strains or species belonging to the genus *Leuconostoc*. In recent days, the whole genome-based taxonomic evaluation became one of the most robust approaches [6]. Our investigation is an attempt to understand the genus *Leuconostoc* whose industrial importance is widely known but have underexplored taxonomic classification.

## Methods

### Genome dataset procurement, initial QC and annotation

All the genomes affiliated to genus *Leuconostoc* available in the public repository as of 31 March 2021 were procured from NCBI (https://www.ncbi.nlm.nih.gov/genome/?term=leuconostoc) and EzBioCloud (https://www.ezbiocloud.net/taxonomy). Procured genomes were quality assessed using checkM v1.1.0 [36] with completeness and contamination bar of 95% and less than 3% respectively. There were altogether 183 including an outgroup genome resources were taken in the study (Table 1). Genome sequences for the type strains of the species *L. lactis* [25], *L. miyukkimchii* [37], *L. palmae* [38], were not included in the analyses due to the absence or poor-quality data in public databases. Also, synonym species/subspecies were strains of the *L. amelibiosum*, *L. argentinum, L. durionis, L. ficulneum*, *L. fructosum*, *L. oeni*, *L. paramesenteroides*, *P. pseudoficulneum* were not included due to their unavailability in the public repository. All the genomes were annotated using prokka v1.12 [39].

**Table 1:** Metadata of the strains used in the study. A detailed description of all the strains used in the study along with the assembly statistics such as assembly size, %GC content, number of CDS, etc. general identification of the strains such as strain Id, NCBI accession number and source etc. are aslo summarized.

| Strains ID | Accession | BioSample | BioProject | Assembly Level | Genome Size(Mb) | GC% | #CDS | #Contigs | #rRNA | #tRNA | #tmRNA | completeness/contamination | Source |
| --- | --- | --- | --- | --- | --- | --- | --- | --- | --- | --- | --- | --- | --- |
| **L. carnosum CBA3620** | GCA_007954525.1 | SAMN11843679 | PRJNA544551 | Complete | 1.70133 | 37.3348 | 1618 | 3 | 12 | 67 | 1 | 99.48/1.05 | kimchi |
| **L. carnosum DSM5576 T** | GCA_014207505.1 | SAMN14908560 | PRJNA632261 | Scaffold | 1.82066 | 37 | 1740 | 16 | 3 | 49 | 1 | 99.48/1.05 | chill-stored meats |
| **L. carnosum JB16** | GCA_000300135.1 | SAMN02603179 | PRJNA174298 | Complete | 1.77347 | 37.091 | 1655 | 5 | 12 | 67 | 1 | 99.48/1.05 | Kimchi, Korean traditional fermented food |
| **L. carnosum MFPA29A1405** | GCA_900322425.1 | SAMEA104699785 | PRJEB25279 | Contig | 1.66684 | 37.2698 | 1598 | 22 | 4 | 48 | 1 | 99.48/1.05 | Beef carpaccio |
| **L. carnosum MFPC16A2803** | GCA_900322415.1 | SAMEA104699786 | PRJEB25280 | Scaffold | 1.81202 | 36.9587 | 1770 | 50 | 3 | 34 | 1 | 99.48/1.05 | Beef carpaccio |
| **L. carnosum WC0318** | GCA_008690985.1 | SAMN11618753 | PRJNA542256 | Scaffold | 1.74508 | 37.2 | 1655 | 11 | 5 | 51 | 1 | 99.48/1.05 | Cooked ham |
| **L. carnosum WC0319** | GCA_008690925.1 | SAMN11618754 | PRJNA542256 | Scaffold | 1.69968 | 37.1 | 1610 | 8 | 5 | 49 | 1 | 99.48/1.05 | Cooked ham |
| **L. carnosum WC0320** | GCA_008690965.1 | SAMN11618755 | PRJNA542256 | Contig | 1.81154 | 37.1 | 1752 | 14 | 5 | 51 | 1 | 99.48/1.05 | Cooked ham |
| **L. carnosum WC0321** | GCA_008690875.1 | SAMN11618756 | PRJNA542256 | Scaffold | 1.80189 | 37.1 | 1735 | 24 | 5 | 52 | 1 | 99.48/1.05 | Cooked ham |
| **L. carnosum WC0322** | GCA_008690865.1 | SAMN11618757 | PRJNA542256 | Scaffold | 1.85285 | 37 | 1806 | 13 | 5 | 50 | 1 | 99.48/1.05 | Cooked ham |
| **L. carnosum WC0323** | GCA_008690845.1 | SAMN11618758 | PRJNA542256 | Scaffold | 1.77288 | 37.2 | 1700 | 17 | 5 | 52 | 1 | 99.48/1.05 | Cooked ham |
| **L. carnosum WC0324** | GCA_008690825.1 | SAMN11618759 | PRJNA542256 | Contig | 1.76563 | 37.1 | 1699 | 12 | 5 | 49 | 1 | 99.48/1.05 | Cooked ham |
| **L. carnosum WC0325** | GCA_008690785.1 | SAMN11618761 | PRJNA542256 | Scaffold | 1.8298 | 37.2 | 1773 | 24 | 5 | 52 | 1 | 99.48/1.05 | Cooked ham |
| **L. carnosum WC0326** | GCA_008689765.1 | SAMN11618763 | PRJNA542256 | Contig | 1.76965 | 37.2 | 1679 | 19 | 5 | 49 | 1 | 99.48/1.05 | Sausage |
| **L. carnosum WC0327** | GCA_008689755.1 | SAMN11618764 | PRJNA542256 | Contig | 1.76938 | 37.2 | 1678 | 18 | 5 | 49 | 1 | 99.48/1.05 | Sausage |
| **L. carnosum WC0328** | GCA_008689695.1 | SAMN11618767 | PRJNA542256 | Scaffold | 1.81458 | 37.1 | 1745 | 27 | 5 | 51 | 1 | 98.95/1.05 | Sausage |
| **L. carnosum WC0329** | GCA_008689675.1 | SAMN11618768 | PRJNA542256 | Scaffold | 1.65083 | 37.2 | 1562 | 13 | 5 | 52 | 1 | 99.48/1.05 | Cooked ham |
| **L. citreum 1301 LGAS** | GCA_001077275.1 | SAMN03197274 | PRJNA267549 | Scaffold | 1.82335 | 39 | 1715 | 37 | 9 | 60 | 1 | 99.89/0.19 | Host: Homo sapiens |
| **L. citreum CBA3621** | GCA_007954785.1 | SAMN11843662 | PRJNA544551 | Complete | 1.90325 | 38.9138 | 1826 | 2 | 12 | 69 | 1 | 99.89/0.56 | kimchi |
| **L. citreum CBA3623** | GCA_007954705.1 | SAMN11843666 | PRJNA544551 | Complete | 1.96941 | 38.9123 | 1848 | 6 | 12 | 69 | 1 | 99.33/0 | kimchi |
| **L. citreum CBA3624** | GCA_007954565.1 | SAMN11843677 | PRJNA544551 | Complete | 1.89477 | 38.9513 | 1778 | 5 | 12 | 69 | 1 | 99.89/0.56 | kimchi |
| **L. citreum CBA3627** | GCA_008033195.1 | SAMN11843672 | PRJNA544551 | Complete | 1.90325 | 38.9138 | 1824 | 2 | 12 | 69 | 1 | 99.89/0.56 | kimchi |
| **L. citreum CW28** | GCA_002591805.1 | SAMN06343424 | PRJNA375706 | Contig | 1.98292 | 38.7 | 1873 | 9 | 12 | 69 | 1 | 99.33/0 | corn fermented beverage |
| **L. citreum DmW 111** | GCA_002115685.1 | SAMN06767174 | PRJNA383560 | Contig | 1.82988 | 38.8 | 1768 | 19 | 7 | 53 | 1 | 99.33/0.75 | from a built environment(Host: Drosophila) |
| **L. citreum EFEL 2700** | GCA_002804045.1 | SAMN08014125 | PRJNA418065 | Chromosome | 1.92383 | 38.964 | 1808 | 5 | 12 | 69 | 1 | 99.33/0 | (host:Brassica rapa subsp. pekinensis) |
| **L. citreum F192-5** | GCA_008326505.1 | SAMD00166981 | PRJDB8200 | Scaffold | 2.06178 | 38.4 | 1989 | 42 | 3 | 47 | 1 | 99.89/0 |  |
| **L. citreum G4** | GCA_006381855.1 | SAMN09099490 | PRJNA470672 | Contig | 1.83552 | 38.8 | 1770 | 4 | 7 | 52 | 1 | 99.89/0 | sourdough |
| **L. citreum KM20** | GCA_000026405.1 | SAMN02603482 | PRJNA16062 | Complete | 1.89661 | 38.8804 | 1777 | 5 | 12 | 69 | 1 | 99.89/0 |  |
| **L. citreum LBAE C10** | GCA_000239895.2 | SAMEA2272656 | PRJEA78993 | Contig | 1.93443 | 38.8 | 1850 | 76 | 3 | 49 | 1 | 99.89/0.08 |  |
| **L. citreum LBAE C11** | GCA_000239915.2 | SAMEA2272136 | PRJEA78995 | Contig | 1.96996 | 38.7 | 1848 | 83 | 3 | 49 | 1 | 99.89/0 |  |
| **L. citreum LBAE E16** | GCA_000239935.2 | SAMEA2272523 | PRJEA78997 | Contig | 1.8016 | 38.9 | 1737 | 45 | 3 | 49 | 1 | 99.89/0 |  |
| **L. citreum NBRC102476** | GCA_007989545.1 | SAMD00166019 | PRJDB6146 | Contig | 1.78757 | 38.9 | 1703 | 25 | 3 | 29 | 1 | 99.89/0 |  |
| **L. citreum NRIC1776 T** | GCA_008326545.1 | SAMD00166982 | PRJDB8200 | Scaffold | 1.80678 | 38.9 | 1737 | 8 | 3 | 47 | 1 | 99.89/0 | Honey dew of rye ear |
| **L. citreum NRRL B-1299** | GCA_000820985.2 | SAMEA2668124 | PRJEB5537 | Scaffold | 1.75381 | 39 | 1769 | 1 | 9 | 53 | 1 | 99.33/0 |  |
| **L. citreum NRRL B-742** | GCA_000820965.2 | SAMEA2668125 | PRJEB5535 | Scaffold | 1.7181 | 39.2 | 1720 | 1 | 9 | 67 | 1 | 99.25/0 |  |
| **L. citreum TMW21194** | GCA_009792915.1 | SAMN07739528 | PRJNA401313 | Contig | 1.73532 | 38.9 | 1639 | 8 | 5 | 51 | 1 | 99.89/0 | food |
| **L. citreum TR116** | GCA_004359915.1 | SAMN07629511 | PRJNA404070 | Contig | 1.83094 | 38.8 | 1764 | 31 | 5 | 51 | 1 | 99.89/0 | yellow pea flour sourdough |
| **L. citreum TR153** | GCA_006406175.1 | SAMN09099486 | PRJNA470672 | Contig | 1.89839 | 38.7 | 1844 | 39 | 5 | 51 | 1 | 99.89/0 | sourdough |
| **L. citreum WiKim0101** | GCA_009707745.1 | SAMN13267039 | PRJNA588925 | Chromosome | 2.01927 | 38.9102 | 1861 | 6 | 12 | 69 | 1 | 99.16/0.19 | kimchi |
| **L. falkenbergense LMG10779 T** | GCA_014634805.1 | SAMD00244137 | PRJDB7793 | Contig | 1.96383 | 39.1 | 1876 | 94 | 3 | 50 | 1 | 100/0.18 | fermentating string beans and traditional yogurt |
| **L. fallax ATCC700006 T** | GCA_004354645.1 | SAMN08555278 | PRJNA434383 | Scaffold | 1.66052 | 37.7 | 1589 | 16 | 15 | 86 | 1 | 99.3/0.52 | sauerkraut |
| **L. gelidum subsp gasicomitatum C120c** | GCA_900009505.1 | SAMEA3718674 | PRJEB12226 | Scaffold | 1.99276 | 36.5 | 1889 | 31 | 3 | 42 | 1 | 99.44/0 | vegetable salad |
| **L. gelidum subsp gasicomitatum C122c** | GCA_900016145.1 | SAMEA3718675 | PRJEB12227 | Scaffold | 2.00836 | 36.6 | 1914 | 22 | 3 | 43 | 1 | 98.88/0.19 | vegetable salad |
| **L. gelidum subsp gasicomitatum KG16-1** | GCA_001536305.1 | SAMEA3608619 | PRJEB11303 | Complete | 2.05073 | 36.8506 | 1963 | 4 | 12 | 67 | 1 | 100/0.11 |  |
| **L. gelidum subsp gasicomitatum KSL4-2** | GCA_900016165.1 | SAMEA3718676 | PRJEB12228 | Scaffold | 2.08949 | 36.6 | 1993 | 23 | 3 | 44 | 1 | 99.33/0.11 | Acetic acid preserved herring with carrots |
| **L. gelidum subsp gasicomitatum LMG18811 T** | GCA_000196855.1 | SAMEA2272393 | PRJEA48123 | Complete | 1.95408 | 36.7 | 1862 | 1 | 12 | 68 | 1 | 99.44/0 | modified-atmosphere packaged, tomato-marinated broiler meat strips |
| **L. gelidum subsp gasicomitatum MFPA44A1401** | GCA_900218135.1 | SAMEA104233143 | PRJEB21908 | Contig | 1.89091 | 36.8 | 1798 | 50 | 2 | 48 | 1 | 100/0 | beef carpaccio |
| **L. gelidum subsp gasicomitatum PB1a** | GCA_900016185.1 | SAMEA3718677 | PRJEB12229 | Scaffold | 2.08363 | 36.6 | 1987 | 29 | 3 | 48 | 1 | 99.44/0.56 | carrot |
| **L. gelidum subsp gasicomitatum PB1e** | GCA_900016175.1 | SAMEA3718678 | PRJEB12230 | Scaffold | 2.08518 | 36.5 | 2014 | 22 | 3 | 43 | 1 | 99.44/0 | carrot |
| **L. gelidum subsp gasicomitatum PL111** | GCA_900016205.1 | SAMEA3718679 | PRJEB12231 | Scaffold | 2.0581 | 36.5 | 1960 | 29 | 3 | 50 | 1 | 99.44/0 | carrot |
| **L. gelidum subsp gasicomitatum TMW 2.1619** | GCA_009296125.1 | SAMN05756392 | PRJNA342276 | Complete | 1.92438 | 36.8534 | 1814 | 4 | 12 | 58 | 1 | 99.64/0.56 | meat |
| **L. gelidum subsp gelidum JB7** | GCA_000298875.1 | SAMN02603178 | PRJNA174297 | Complete | 1.8935 | 36.7 | 1768 | 1 | 12 | 67 | 1 | 99.44/0.19 | kimchi, Korean traditional fermented food |
| **L. gelidum subsp gelidum KCTC3527 T** | GCA_000166715.2 | SAMN02470198 | PRJNA60229 | Contig | 1.95728 | 36.6 | 1845 | 43 | 3 | 47 | 1 | 99.44/0.19 | vacuum-packed meats |
| **L. holzapfelii CCUG54536 T** | GCA_012396485.1 | SAMN14517846 | PRJNA622446 | Contig | 1.75843 | 44 | 1668 | 18 | 5 | 50 | 1 | 99.37/0.28 | coffee fermentation |
| **L. inhae KCTC3774 T** | GCA_000166735.2 | SAMN02472184 | PRJNA60231 | Contig | 2.29809 | 36.4 | 1622 | 893 | 3 | 48 | 1 | 95.81/6.07 | kimchi |
| **L. kimchii C2** | GCA_000219785.1 | SAMN02603135 | PRJNA67911 | Complete | 1.87727 | 37.9 | 1792 | 1 | 12 | 68 | 1 | 100/0 | kimchi |
| **L. kimchii IMSNU11154 T** | GCA_000092505.1 | SAMN02603394 | PRJNA40837 | Complete | 2.10179 | 37.9101 | 2025 | 6 | 12 | 68 | 1 | 100/0 | kimchi, a traditional Korean food |
| **L. kimchii NKJ218** | GCA_004551615.1 | SAMN10887832 | PRJNA521423 | Complete | 1.99344 | 37.7797 | 1892 | 4 | 12 | 68 | 1 | 100/0 | kimchi |
| **L. lactis 1300 LCIT** | GCA_001062635.1 | SAMN03197273 | PRJNA267549 | Scaffold | 1.80711 | 43.1 | 1687 | 58 | 9 | 63 | 1 | 99.44/0.26 | Host: Homo sapiens |
| **L. lactis aa 0143** | GCA_004167235.1 | SAMN10239554 | PRJNA496358 | Contig | 1.73814 | 43.2 | 1637 | 38 | 3 | 46 | 1 | 99.44/0 | stool(host:Homo sapiens) |
| **L. lactis BIOML-A1** | GCA_009678855.1 | SAMN11946164 | PRJNA544527 | Contig | 1.748 | 43.1 | 1632 | 30 | 4 | 46 | 1 | 99.44/0 | (host:Homo sapiens) |
| **L. lactis CBA3625** | GCA_007954605.1 | SAMN11843673 | PRJNA544551 | Complete | 1.79161 | 43.3467 | 1714 | 2 | 12 | 68 | 1 | 99.44/0.56 | kimchi |
| **L. lactis CBA3626** | GCA_007954665.1 | SAMN11843668 | PRJNA544551 | Complete | 1.83981 | 43.1491 | 1742 | 2 | 12 | 70 | 1 | 99.44/1.12 | kimchi |
| **L. lactis CCK940** | GCA_002287365.1 | SAMN07482638 | PRJNA397634 | Contig | 1.74151 | 43.3 | 1657 | 2 | 12 | 69 | 1 | 100/0 | kimchi |
| **L. lactis KACC91922** | GCA_000709265.1 | SAMN02730106 | PRJNA244961 | Contig | 1.68817 | 43.4 | 1619 | 35 | 3 | 50 | 1 | 99.44/0.67 | Kimchi |
| **L. lactis KCTC3773** | GCA_000179875.1 | SAMN02470199 | PRJNA52539 | Contig | 1.72068 | 42.9 | 1663 | 98 | 3 | 48 | 1 | 99.44/0 |  |
| **L. lactis KFRI01** | GCA_001998805.1 | SAMN05219201 | PRJNA325075 | Complete | 1.71519 | 43.3281 | 1613 | 3 | 12 | 69 | 1 | 99.44/0 | kimchi |
| **L. lactis LN19** | GCA_002092595.1 | SAMN05981500 | PRJNA352459 | Contig | 1.72439 | 42.9 | 1633 | 47 | 6 | 51 | 1 | 99.44/0.14 | dairy |
| **L. lactis LN24** | GCA_002092695.1 | SAMN05981502 | PRJNA352459 | Contig | 1.72466 | 42.9 | 1641 | 48 | 6 | 52 | 1 | 99.44/0.14 | dairy |
| **L. lactis NBRC 12455** | GCA_006539105.1 | SAMD00097163 | PRJDB6024 | Contig | 1.64349 | 43.5 | 1531 | 49 | 3 | 39 |  | 99.44/0 |  |
| **L. lactis WiKim40** | GCA_001698145.1 | SAMN05300424 | PRJNA327015 | Complete | 1.78807 | 43.1131 | 1654 | 4 | 12 | 68 | 1 | 100/0.56 | Kimchi |
| **L. litichii MB7 T** | GCA_008107645.1 | SAMN10786216 | PRJNA516297 | Scaffold | 1.86586 | 35.8 | 1789 | 13 | 3 | 48 | 1 | 100/0 | Litchi chinensis |
| **L. mesenteroides CBA3628** | GCA_007954745.1 | SAMN11843664 | PRJNA544551 | Complete | 1.86456 | 36.9975 | 1757 | 4 | 12 | 71 | 1 | 100/0 | kimchi |
| **L. mesenteroides subsp cremoris ATCC19254 T** | GCA_000160595.1 | SAMN00139187 | PRJNA34629 | Scaffold | 1.73507 | 38.5 | 1511 | 29 | 3 | 49 | 1 | 100/0 | Hansen's dried starter powder |
| **L. mesenteroides**<br>**subsp cremoris**<br>**LMGCF01** | GCA_002092045.1 | SAMN05981463 | PRJNA352459 | Contig | 1.64067 | 38 | 1449 | 115 | 5 | 51 | 1 | 99.82/0 | dairy |
| **L. mesenteroides**<br>**subsp cremoris**<br>**LMGCF02** | GCA_002092135.1 | SAMN05981464 | PRJNA352459 | Contig | 1.72228 | 37.9 | 1570 | 112 | 5 | 53 | 1 | 100/0 | dairy |
| **L. mesenteroides**<br>**subsp cremoris**<br>**LMGCF03** | GCA_002092145.1 | SAMN05981465 | PRJNA352459 | Contig | 1.66544 | 38 | 1494 | 116 | 5 | 53 | 1 | 100/0 | dairy |
| **L. mesenteroides**<br>**subsp cremoris**<br>**LMGCF04** | GCA_002092155.1 | SAMN05981466 | PRJNA352459 | Contig | 1.64019 | 38 | 1446 | 124 | 5 | 52 | 1 | 100/0 | dairy |
| **L. mesenteroides**<br>**subsp cremoris**<br>**LMGCF05** | GCA_002092085.1 | SAMN05981467 | PRJNA352459 | Contig | 1.63731 | 38 | 1452 | 118 | 5 | 53 | 1 | 100/0 | dairy |
| **L. mesenteroides**<br>**subsp cremoris**<br>**LMGCF07** | GCA_002092215.1 | SAMN05981469 | PRJNA352459 | Scaffold | 1.63929 | 38.1 | 1431 | 109 | 5 | 53 | 1 | 100/0 | dairy |
| **L. mesenteroides**<br>**subsp cremoris**<br>**LMGCF09** | GCA_002092105.1 | SAMN05981471 | PRJNA352459 | Contig | 1.62909 | 38.1 | 1468 | 105 | 5 | 53 | 1 | 100/0 | dairy |
| **L. mesenteroides**<br>**subsp cremoris**<br>**LMGCF10** | GCA_002092245.1 | SAMN05981472 | PRJNA352459 | Contig | 1.61642 | 38 | 1452 | 109 | 5 | 51 | 1 | 100/0 | dairy |
| **L. mesenteroides**<br>**subsp cremoris**<br>**LMGCF11** | GCA_002092315.1 | SAMN05981473 | PRJNA352459 | Contig | 1.67437 | 38.2 | 1488 | 86 | 5 | 53 | 1 | 99.87/0 | dairy |
| **L. mesenteroides**<br>**subsp cremoris**<br>**LMGCF12** | GCA_002092385.1 | SAMN05981474 | PRJNA352459 | Contig | 1.66343 | 38.1 | 1474 | 113 | 4 | 54 | 1 | 100/0.53 | dairy |
| **L. mesenteroides**<br>**subsp cremoris**<br>**LMGCF13** | GCA_002092465.1 | SAMN05981475 | PRJNA352459 | Contig | 1.63644 | 38.3 | 1451 | 219 | 4 | 50 | 1 | 100/1.06 | dairy |
| **L. mesenteroides**<br>**subsp cremoris**<br>**LMGCF14** | GCA_002092335.1 | SAMN05981476 | PRJNA352459 | Contig | 1.62752 | 38 | 1462 | 105 | 5 | 52 | 1 | 100/0 | dairy |
| **L. mesenteroides**<br>**subsp cremoris**<br>**LMGCF16** | GCA_002092515.1 | SAMN05981478 | PRJNA352459 | Contig | 1.61323 | 38 | 1448 | 124 | 5 | 52 | 1 | 100/0 | dairy |
| **L. mesenteroides**<br>**subsp cremoris**<br>**LMGCF17** | GCA_002092425.1 | SAMN05981479 | PRJNA352459 | Contig | 1.62949 | 38 | 1457 | 101 | 5 | 52 | 1 | 100/0 | dairy |
| **L. mesenteroides**<br>**subsp cremoris**<br>**LMGCF18** | GCA_002092495.1 | SAMN05981480 | PRJNA352459 | Contig | 1.62399 | 38 | 1456 | 105 | 5 | 52 | 1 | 100/0 | dairy |
| **L. mesenteroides subsp cremoris LMGCF19** | GCA_002092415.1 | SAMN05981481 | PRJNA352459 | Contig | 1.64533 | 38 | 1517 | 113 | 5 | 53 | 1 | 100/0 | dairy |
| **L. mesenteroides subsp cremoris LMGCF20** | GCA_002092455.1 | SAMN05981482 | PRJNA352459 | Contig | 1.63087 | 38 | 1474 | 109 | 5 | 53 | 1 | 99.74/0 | dairy |
| **L. mesenteroides subsp cremoris LN07** | GCA_002092615.1 | SAMN05981497 | PRJNA352459 | Contig | 1.59621 | 38.1 | 1424 | 107 | 5 | 52 | 1 | 100/0 | dairy |
| **L. mesenteroides subsp cremoris T26** | GCA_000686485.1 | SAMN02767914 | PRJNA205545 | Scaffold | 1.83247 | 38.4 | 1690 | 123 | 3 | 56 | 1 | 100/10.23 | mesophilic cheese starter cultures |
| **L. mesenteroides subsp cremoris TIFN8** | GCA_000447945.1 | SAMN02472108 | PRJNA175676 | Contig | 1.71009 | 38.2 | 1452 | 173 | 7 | 61 |  | 99.21/0.26 | propagated starter culture used in cheese manufacturing |
| **L. mesenteroides subsp dextranicum DSM20484 T** | GCA_001047695.1 | SAMN03105774 | PRJNA263697 | Complete | 1.85473 | 38.0389 | 1692 | 2 | 12 | 71 | 1 | 100/0.79 | Cheese |
| **L. mesenteroides subsp dextranicum FAM 18356** | GCA_005864365.1 | SAMN11653949 | PRJNA543085 | Contig | 1.87153 | 37.8 | 1707 | 40 | 5 | 53 | 1 | 100/0 |  |
| **L. mesenteroides subsp dextranicum FM06** | GCA_002148235.1 | SAMN06061940 | PRJNA355067 | Complete | 1.94688 | 37.9649 | 1746 | 4 | 12 | 71 | 1 | 100/0 | 10 weeks old 45+ Samso cheese |
| **L. mesenteroides subsp dextranicum LbT16** | GCA_001184265.1 | SAMN03463677 | PRJNA280520 | Contig | 1.90636 | 37.8 | 1733 | 65 | 3 | 51 | 1 | 99.47/0 | Taleggio cheese |
| **L. mesenteroides subsp dextranicum LN05** | GCA_002092565.1 | SAMN05981496 | PRJNA352459 | Contig | 2.00187 | 38 | 1835 | 250 | 16 | 57 | 1 | 98.53/2.95 | dairy |
| **L. mesenteroides subsp dextranicum LN08** | GCA_002092625.1 | SAMN05981498 | PRJNA352459 | Contig | 1.96519 | 37.8 | 1866 | 106 | 5 | 54 | 1 | 100/0 | dairy |
| **L. mesenteroides subsp dextranicum LN32** | GCA_002092705.1 | SAMN05981505 | PRJNA352459 | Contig | 1.84828 | 37.8 | 1696 | 68 | 5 | 54 | 1 | 100/0 | dairy |
| **L. mesenteroides subsp dextranicum LN34** | GCA_002092775.1 | SAMN05981506 | PRJNA352459 | Contig | 1.87513 | 37.7 | 1719 | 110 | 5 | 53 | 1 | 100/0 | dairy |
| **L. mesenteroides subsp dextranicum NCTC10817** | GCA_900452955.1 | SAMEA4362418 | PRJEB6403 | Contig | 1.97325 | 37.8 | 1860 | 3 | 12 | 71 | 1 | 100/0.53 |  |
| **L. mesenteroides subsp dextranicum OG2** | GCA_002276945.1 | SAMN06718482 | PRJNA375758 | Contig | 1.71628 | 37.9 | 1583 | 146 | 7 | 51 | 1 | 100/0 | kefir |
| **L. mesenteroides subsp dextranicum P45** | GCA_000756355.1 | SAMN03032191 | PRJNA260860 | Contig | 1.87419 | 37.5 | 1726 | 6 | 9 | 52 | 1 | 100/0 | fermented pulque |
| L. mesenteroides<br>subsp dextranicum<br>WC0331 | GCA_008690745.1 | SAMN11618762 | PRJNA542256 | Scaffold | 2.01813 | 37.6 | 1923 | 10 | 5 | 53 | 1 | 100/0.53 | Cooked ham |
| L. mesenteroides<br>subsp<br>jonggajibkimchii B5 | GCA_004745755.1 | SAMN10484942 | PRJNA507123 | Contig | 2.12682 | 37.5 | 2037 | 26 | 5 | 53 | 1 | 100/0.18 | Mike cake |
| L. mesenteroides<br>subsp<br>jonggajibkimchii<br>BD3749 | GCA_001583825.1 | SAMN04508169 | PRJNA312878 | Complete | 1.98657 | 37.8 | 1898 | 1 | 12 | 71 | 1 | 100/0 | fermeted vegetable |
| L. mesenteroides<br>subsp<br>jonggajibkimchii<br>CBA7131 | GCA_003255835.1 | SAMN07206913 | PRJNA390217 | Complete | 2.08731 | 37.6375 | 1986 | 4 | 12 | 71 | 1 | 99.47/0 | Korean adult feces |
| L. mesenteroides<br>subsp<br>jonggajibkimchii<br>DRC1506 T | GCA_001886915.1 | SAMN04198570 | PRJNA299366 | Complete | 1.98392 | 37.6424 | 1787 | 4 | 12 | 71 | 1 | 99.89/0.53 | Kimchi |
| L. mesenteroides<br>subsp<br>jonggajibkimchii KFRI-MG | GCA_000512955.1 | SAMN02641526 | PRJNA18679 | Complete | 1.89641 | 37.7 | 1796 | 1 | 12 | 69 | 1 | 100/0 |  |
| L. mesenteroides<br>subsp<br>jonggajibkimchii<br>KIBGE-IB22 | GCA_005049065.1 | SAMN11431368 | PRJNA532993 | Scaffold | 1.94823 | 37.5 | 1880 | 13 | 3 | 50 | 1 | 100/0.18 | cabbage(host:Vegetables) |
| L. mesenteroides<br>subsp<br>jonggajibkimchii<br>SRCM102735 | GCA_009913935.1 | SAMN08707610 | PRJNA438180 | Complete | 2.05844 | 37.5883 | 1982 | 4 | 12 | 71 | 1 | 100/0 | Soybean paste (Chonggugjang) |
| L. mesenteroides<br>subsp<br>jonggajibkimchii<br>SRCM103356 | GCA_004102585.1 | SAMN10742283 | PRJNA515114 | Complete | 2.03939 | 37.5826 | 1984 | 4 | 12 | 71 | 1 | 100/0.18 | food |
| L. mesenteroides<br>subsp<br>jonggajibkimchii<br>SRCM103453 | GCA_004103675.1 | SAMN10754493 | PRJNA515383 | Complete | 2.1826 | 37.6453 | 2090 | 5 | 12 | 71 | 1 | 100/0 | food |
| L. mesenteroides<br>subsp<br>jonggajibkimchii<br>SRCM103460 | GCA_004194375.1 | SAMN10743321 | PRJNA515149 | Complete | 2.08721 | 37.5882 | 1996 | 5 | 12 | 71 | 1 | 100/0 | food |
| L. mesenteroides<br>subsp<br>jonggajibkimchii<br>WC0333 | GCA_008689685.1 | SAMN11618766 | PRJNA542256 | Contig | 2.08859 | 37.5 | 2031 | 35 | 5 | 53 | 1 | 100/0 | Sausage |
| **L. mesenteroides**<br>subsp<br>jonggajibkimchii<br>WiKim33 | GCA_003433375.1 | SAMN07156024 | PRJNA387486 | Complete | 1.97006 | 37.6362 | 1883 | 4 | 12 | 71 | 1 | 100/0 | Baechu-kimchi |
| **L. mesenteroides**<br>subsp mesenteroides<br>CBA3607 | GCA_009676745.1 | SAMN13258970 | PRJNA544551 | Complete | 1.85432 | 37.008 | 1750 | 3 | 12 | 71 | 1 | 100/0 | kimchi |
| **L. mesenteroides**<br>subsp mesenteroides<br>213M0 | GCA_001570665.1 | SAMD00042708 | PRJDB4328 | Contig | 2.03285 | 37.7 | 1946 | 58 | 3 | 49 | 1 | 99.93/0.53 |  |
| **L. mesenteroides**<br>subsp mesenteroides<br>406 | GCA_001515385.1 | SAMD00042709 | PRJDB4329 | Contig | 2.00431 | 37.7 | 1932 | 69 | 3 | 49 | 1 | 100/1.06 |  |
| **L. mesenteroides**<br>subsp mesenteroides<br>ATCC8293 T | GCA_000014445.1 | SAMN02598482 | PRJNA315 | Complete | 2.07576 | 37.6586 | 1967 | 2 | 12 | 71 | 1 | 100/0.53 | fermenting olives |
| **L. mesenteroides**<br>subsp mesenteroides<br>AtHG50 | GCA_004368765.1 | SAMN08777814 | PRJNA441539 | Scaffold | 2.03285 | 37.7 | 1960 | 60 | 3 | 52 | 1 | 100/0.53 |  |
| **L. mesenteroides**<br>subsp mesenteroides<br>BD1710 | GCA_002117185.1 | SAMN04530070 | PRJNA314170 | Complete | 2.12404 | 37.6 | 2014 | 1 | 12 | 71 | 1 | 100/0.53 | kefir |
| **L. mesenteroides**<br>subsp mesenteroides<br>CECT9217 | GCA_900290445.1 | SAMEA104569583 | PRJEB24702 | Contig | 2.02219 | 37.7 | 1932 | 13 | 5 | 53 | 1 | 100/0 |  |
| **L. mesenteroides**<br>subsp mesenteroides<br>CECT9266 | GCA_900312985.1 | SAMEA104569776 | PRJEB24710 | Contig | 2.10613 | 37.7 | 2032 | 44 | 5 | 53 | 1 | 99.82/0 |  |
| **L. mesenteroides**<br>subsp mesenteroides<br>CECT9268 | GCA_900290155.1 | SAMEA104569777 | PRJEB24712 | Contig | 2.07557 | 37.7 | 2002 | 26 | 5 | 53 | 1 | 99.82/0 |  |
| **L. mesenteroides**<br>subsp mesenteroides<br>DE0394 | GCA_007673005.1 | SAMN11792554 | PRJNA543692 | Scaffold | 1.93918 | 37.7 | 1816 | 358 | 3 | 48 | 1 | 100/1.6 | environmental |
| **L. mesenteroides**<br>subsp mesenteroides<br>DRC0211 | GCA_002009375.1 | SAMN04198273 | PRJNA299363 | Complete | 2.1241 | 37.5981 | 1962 | 5 | 12 | 71 | 1 | 100/0 | Kimchi |
| **L. mesenteroides**<br>subsp mesenteroides<br>GL1 | GCA_001541125.1 | SAMN04231183 | PRJNA300958 | Scaffold | 1.81597 | 38.1 | 1649 | 11 | 2 | 35 | 1 | 100/0.79 | Dromedary milk |
| **L. mesenteroides**<br>subsp mesenteroides<br>J18 | GCA_000234825.3 | SAMN02603136 | PRJNA75083 | Complete | 2.01726 | 37.6784 | 1883 | 5 | 12 | 71 | 1 | 100/0 |  |
| **L. mesenteroides**<br>subsp mesenteroides<br>KMB608 | GCA_003346165.1 | SAMN09398940 | PRJNA474823 | Contig | 1.95951 | 37.7 | 1877 | 81 | 3 | 46 | 1 | 100/0 | sheep milk |
| **L. mesenteroides**<br>subsp **mesenteroides**<br>**KMB609** | GCA_003346155.1 | SAMN09398941 | PRJNA474823 | Contig | 2.01223 | 37.7 | 1930 | 84 | 3 | 47 | 1 | 100/0.53 | bryndza cheese |
| **L. mesenteroides**<br>subsp **mesenteroides**<br>**KMB611** | GCA_003346345.1 | SAMN09398943 | PRJNA474823 | Contig | 2.10341 | 37.5 | 2037 | 33 | 3 | 46 | 1 | 100/0.53 | bryndza cheese |
| **L. mesenteroides**<br>subsp **mesenteroides**<br>**LbE15** | GCA_001184245.1 | SAMN03463642 | PRJNA280507 | Contig | 2.00792 | 37.6 | 1923 | 63 | 3 | 51 | 1 | 100/0 | Taleggio cheese |
| **L. mesenteroides**<br>subsp **mesenteroides**<br>**LbE16** | GCA_001184255.1 | SAMN03463665 | PRJNA280516 | Contig | 2.03609 | 37.5 | 1964 | 85 | 3 | 51 | 1 | 100/0.53 | Taleggio cheese |
| **L. mesenteroides**<br>subsp **mesenteroides**<br>**LK-151** | GCA_002370415.1 | SAMD00065790 | PRJDB5233 | Complete | 2.13737 | 37.7366 | 2017 | 4 | 9 | 68 | 1 | 100/0 |  |
| **L. mesenteroides**<br>subsp **mesenteroides**<br>**LN25** | GCA_002092735.1 | SAMN05981503 | PRJNA352459 | Contig | 2.2776 | 37.5 | 2180 | 70 | 6 | 54 | 1 | 100/1.06 | dairy |
| **L. mesenteroides**<br>subsp **mesenteroides**<br>**LN27** | GCA_002092745.1 | SAMN05981504 | PRJNA352459 | Contig | 2.2246 | 37.5 | 2123 | 54 | 6 | 54 | 1 | 100/1.06 | dairy |
| **L. mesenteroides**<br>subsp **mesenteroides**<br>**M11** | GCA_009497095.1 | SAMN13018494 | PRJNA577063 | Scaffold | 1.97939 | 37.8 | 1863 | 79 | 3 | 52 | 1 | 100/1.06 | (host:Oreochromis niloticus) |
| **L. mesenteroides**<br>subsp **mesenteroides**<br>**NBRC102481** | GCA_007989505.1 | SAMD00166021 | PRJDB6187 | Contig | 1.69543 | 37.8 | 1630 | 109 | 3 | 45 | 1 | 98.41/0 |  |
| **L. mesenteroides**<br>subsp **mesenteroides**<br>**NBRC3832** | GCA_006538765.1 | SAMD00093693 | PRJDB6025 | Contig | 1.90981 | 37.6 | 1789 | 31 | 3 | 45 | 1 | 100/0.53 |  |
| **L. mesenteroides**<br>subsp **mesenteroides**<br>**SRCM102733** | GCA_009913915.1 | SAMN08707608 | PRJNA438180 | Complete | 2.05873 | 37.7211 | 1967 | 3 | 12 | 71 | 1 | 100/0 | Gochujang |
| **L. mesenteroides**<br>subsp **mesenteroides**<br>**TR154** | GCA_006406185.1 | SAMN09099487 | PRJNA470672 | Contig | 1.85995 | 37.8 | 1756 | 43 | 5 | 53 | 1 | 100/0 | sourdough |
| **L. mesenteroides**<br>subsp **mesenteroides**<br>**WC0330** | GCA_008690805.1 | SAMN11618760 | PRJNA542256 | Scaffold | 1.76569 | 37.9 | 1637 | 41 | 4 | 54 | 1 | 100/0.26 | Cooked ham |
| **L. mesenteroides**<br>subsp **mesenteroides**<br>**WC0332** | GCA_008689715.1 | SAMN11618765 | PRJNA542256 | Contig | 1.8553 | 37.7 | 1744 | 39 | 5 | 54 | 1 | 100/0 | Sausage |
| **L. mesenteroides**<br>subsp **mesenteroides**<br>**Wikim17** | GCA_000787735.1 | SAMD00022975 | PRJDB3305 | Contig | 1.85932 | 37.8 | 1748 | 41 | 4 | 53 | 1 | 100/0.26 |  |
| **L. mesenteroides subsp mesenteroides YL48** | GCA_002886025.1 | SAMN06298812 | PRJNA371414 | Contig | 2.0584 | 37.6 | 1975 | 154 | 6 | 54 | 1 | 100/0 | Root Surface(host:Carrot) |
| **L. pseudomesenteroides 1159** | GCA_000686465.1 | SAMN02767913 | PRJNA205546 | Scaffold | 2.03869 | 39 | 1969 | 100 | 3 | 49 | 1 | 100/0 | mesophilic cheese starter cultures |
| **L. pseudomesenteroides 4882** | GCA_000297375.1 | SAMEA3138421 | PRJNA84285 | Contig | 2.00859 | 39.1 | 1936 | 106 | 5 | 59 | 1 | 93.12/0.18 |  |
| **L. pseudomesenteroides AMBR10** | GCA_901830415.1 | SAMEA5665507 | PRJEB32716 | Contig | 2.29673 | 39.2 | 2248 | 338 | 7 | 56 | 1 | 100/0.7 | Human adenoid |
| **L. pseudomesenteroides BM2** | GCA_002092535.1 | SAMN05981461 | PRJNA352459 | Contig | 2.02255 | 39 | 1997 | 101 | 7 | 53 | 1 | 100/0 | dairy |
| **L. pseudomesenteroides CBA3630** | GCA_008033175.1 | SAMN11843675 | PRJNA544551 | Complete | 2.31609 | 38.9999 | 2171 | 4 | 12 | 70 | 1 | 99.82/0.18 | kimchi |
| **L. pseudomesenteroides HPK01** | GCA_002092075.1 | SAMN05981330 | PRJNA352459 | Contig | 1.95326 | 39.1 | 1853 | 92 | 5 | 51 | 1 | 100/0 | dairy starter |
| **L. pseudomesenteroides KMB610** | GCA_003346375.1 | SAMN09398942 | PRJNA474823 | Contig | 1.9824 | 39 | 1919 | 57 | 3 | 43 | 1 | 100/0 | bryndza cheese |
| **L. pseudomesenteroides LMG11482 T** | GCA_014634745.1 | SAMD00239684 | PRJDB7793 | Contig | 2.09581 | 38.9 | 1971 | 29 | 4 | 50 | 1 | 99.82/0 | cane juice |
| **L. pseudomesenteroides LMGCF06** | GCA_002092035.1 | SAMN05981468 | PRJNA352459 | Contig | 1.95205 | 39.1 | 1848 | 83 | 5 | 51 | 1 | 100/0 | dairy |
| **L. pseudomesenteroides LMGCF08** | GCA_002092165.1 | SAMN05981470 | PRJNA352459 | Contig | 2.19336 | 39.1 | 2069 | 267 | 5 | 53 | 1 | 100/11.2 | dairy |
| **L. pseudomesenteroides LMGCF15** | GCA_002092375.1 | SAMN05981477 | PRJNA352459 | Contig | 1.92427 | 39.1 | 1833 | 78 | 5 | 51 | 1 | 100/0 | dairy |
| **L. pseudomesenteroides LMGH100** | GCA_002072505.1 | SAMN05981487 | PRJNA352459 | Contig | 2.06006 | 39.1 | 1903 | 238 | 4 | 52 | 1 | 98.7/2.61 | dairy |
| **L. pseudomesenteroides LMGH278** | GCA_002072495.1 | SAMN05981488 | PRJNA352459 | Contig | 1.9401 | 39.1 | 1852 | 106 | 5 | 51 | 1 | 100/0 | dairy |
| **L. pseudomesenteroides LMGH280** | GCA_002072555.1 | SAMN05981489 | PRJNA352459 | Contig | 1.98015 | 39.1 | 1861 | 85 | 4 | 55 | 1 | 100/0.71 | dairy |
| **L. pseudomesenteroides LMGH284** | GCA_002072575.1 | SAMN05981490 | PRJNA352459 | Scaffold | 2.07199 | 39 | 1952 | 222 | 4 | 52 | 1 | 99.21/2.03 | dairy |
| **L. pseudomesenteroides LMGH61** | GCA_002072515.1 | SAMN05981483 | PRJNA352459 | Scaffold | 2.00878 | 39.1 | 1901 | 62 | 4 | 54 | 1 | 100/0 | dairy |
| **L. pseudomesenteroides LMGH83** | GCA_002072475.1 | SAMN05981484 | PRJNA352459 | Contig | 2.05883 | 39.1 | 1885 | 201 | 4 | 53 | 1 | 98.4/1.41 | dairy |
| **L. pseudomesenteroides LMGH95** | GCA_002072565.1 | SAMN05981485 | PRJNA352459 | Contig | 1.98982 | 39 | 1948 | 171 | 5 | 51 | 1 | 100/0.18 | dairy |
| **L. pseudomesenteroides LMGH97** | GCA_002072585.1 | SAMN05981486 | PRJNA352459 | Contig | 1.98975 | 39 | 1946 | 132 | 5 | 50 | 1 | 100/0 | dairy |
| **L. pseudomesenteroides LMGTW1** | GCA_002092235.1 | SAMN05981491 | PRJNA352459 | Contig | 1.9692 | 39 | 1924 | 125 | 5 | 52 | 1 | 100/0 | dairy |
| **L. pseudomesenteroides LMGTW3** | GCA_002092355.1 | SAMN05981492 | PRJNA352459 | Contig | 1.89805 | 39.1 | 1834 | 87 | 5 | 52 | 1 | 100/0 | dairy |
| **L. pseudomesenteroides LMGTW6** | GCA_002092255.1 | SAMN05981493 | PRJNA352459 | Contig | 1.88144 | 39.1 | 1821 | 90 | 5 | 52 | 1 | 100/0 | dairy |
| **L. pseudomesenteroides LMGTW8** | GCA_002092295.1 | SAMN05981494 | PRJNA352459 | Contig | 1.98207 | 39 | 1936 | 118 | 5 | 52 | 1 | 100/0 | dairy |
| **L. pseudomesenteroides LN02** | GCA_002092555.1 | SAMN05981495 | PRJNA352459 | Contig | 1.91243 | 39.1 | 1803 | 88 | 6 | 53 | 1 | 100/0 | dairy |
| **L. pseudomesenteroides LN12** | GCA_002092635.1 | SAMN05981499 | PRJNA352459 | Contig | 1.86409 | 39.1 | 1762 | 85 | 6 | 52 | 1 | 100/0 | dairy |
| **L. pseudomesenteroides LN23** | GCA_002092645.1 | SAMN05981501 | PRJNA352459 | Contig | 1.99273 | 39.1 | 1936 | 126 | 5 | 47 | 1 | 100/0.18 | dairy |
| **L. pseudomesenteroides MGYG-HGUT-00096** | GCA_902363165.1 | SAMEA5849597 | PRJEB33885 | Scaffold | 2.29682 |  |  | 33 | 4 | 52 | 1 | 99.82/0 | human gut |
| **L. pseudomesenteroides PS12** | GCA_000686505.1 | SAMN02767912 | PRJNA205547 | Scaffold | 1.93438 | 39.1 | 1863 | 91 | 3 | 48 | 1 | 100/0 | undefined mesophilic cheese starter culture |
| **L. pseudomesenteroides TR070** | GCA_006382035.1 | SAMN09099485 | PRJNA470672 | Contig | 2.23055 | 38.8 | 2162 | 76 | 5 | 52 | 1 | 99.82/0 | sourdough |
| **L. rapi DSM27776 T** | GCA_016908715.1 | SAMN17619917 | PRJNA695649 | Contig | 1.92491 | 38 | 1843 | 24 | 3 | 49 | 1 | 100/1.24 | rutabaga |
| **L. suionicum CECT 8486** | GCA_900289255.1 | SAMEA104569300 | PRJEB24691 | Contig | 2.07002 | 37.3 | 1968 | 20 | 5 | 53 | 1 | 100/0.53 |  |
| **L. suionicum CECT 9216** | GCA_900290145.1 | SAMEA104569335 | PRJEB24693 | Contig | 2.06953 | 37.3 | 1972 | 18 | 5 | 53 | 1 | 100/0.53 |  |
| **L. suionicum CECT8484** | GCA_900289205.1 | SAMEA104569299 | PRJEB24690 | Contig | 2.06903 | 37.3 | 1974 | 18 | 5 | 53 | 1 | 100/0.53 |  |
| **L. suionicum DSM20241 T** | GCA_001891125.1 | SAMN04811211 | PRJNA318320 | Complete | 2.04883 | 37.5936 | 1923 | 2 | 12 | 71 | 1 | 100/0 | Source unknown |

### Phylogenetic and Phylogenomic analysis

16S rRNA gene predicted using barrnap v0.8 (https://github.com/tseemann/barrnap) was used to fetch 16S rRNA gene sequences from all 183 genomes. Multiple sequence alignment of 16S rRNA was performed using clustalw [40]. A phylogenetic tree based on maximum likelihood (ML) was constructed using mega 7 [41] with a bootstrap replication of 1000. *Weissella viridescens* DSM 20410 was used as an outgroup. Whole genome-based phylogeny was implemented using PhyloPhlAn v3.0 [42] which used more than 400 most conserved genes across the strains under investigation. PhyloPhlAn uses usearch [43] for searching the gene cluster, muscle [44] for performing multiple sequence alignment and fasttree [45] for phylogenetic tree generation. A core genome-based tree was also generated using roary v3.1.2 [46] with 70% (inter-species similarity) cut off from the core genome alignment obtained with mafft [47]. All the phylogenetic tree generated was annotated using iTOL (https://itol.embl.de/) [48] for more lucid representation.

### Genome similarity assessment

Genome similarity assessment was performed using several methods. Average nucleotide identity (ANI) was calculated using orthoANI v1.2 [49, 50] which uses usearch v8.1 [43] to find the orthologous cluster across the genomes for similarity comparison. A genome similarity assessment cut-off of 96% for orthoANI was used. Another tool fastANI v1.32 [51] (https://github.com/ParBLiSS/FastANI) was implemented to calculate nucleotide identity. fastANI uses a novel algorithm, which utilizes Mashmap with an intra and inter-species demarcation of >96% and <83% [52]. Digital DNA-DNA hybridization (dDDH) [53] was implemented using an online web portal of genome-to-genome distance calculator 2.1 (GGDC) (http://ggdc.dsmz.de/ggdc.php#). Heatmap of the genome similarity assessment was generated using GENE-E (https://software.broadinstitute.org/GENE-E/).

### Pangenome and core genome assessment

Pangenome analysis was implemented using roary v3.1.2 [46] with gff files obtained from prokka with blastp percentage identity of 85%. The flower plot indicates the core gene, unique gene and species specific to genes (Figure 3A). fasttree v2.1.10 [45] was implemented on the core gene alignment to obtain the core gene phylogeny. Core genome obtained was annotated with eggNOG-mapper [54] (http://eggnog-mapper.embl.de/).

**Figure 1:**
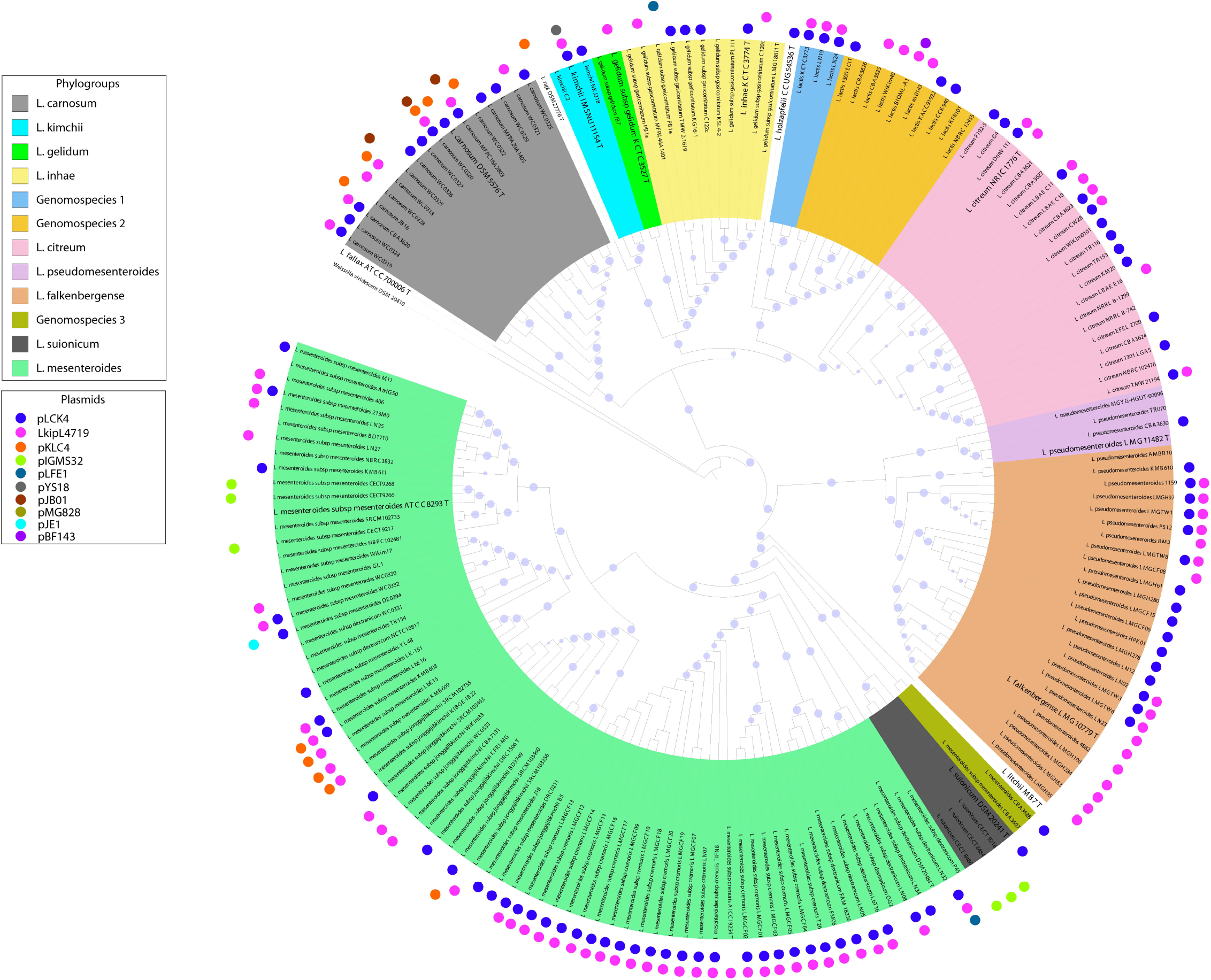
Whole genome-based core genome phylogenomic tree obtained using roary: Phylogenetic tree was obtained using fasttree on the core gene alignment using mafft in roary run. For circular representation iTOL was used and labelled in accordance to the species group. Bootstrap values are represented with blue dots. *Weissella viridescens* DSM 20410 was used as an outgroup. Genomospecies 1, genomospecies 2 and genomospecies 3 represent the novel group of bacteria within the genus *Leuconostoc*. The plasmid was labelled in the genome with coloured dots at the circumference.

**Figure 2:**
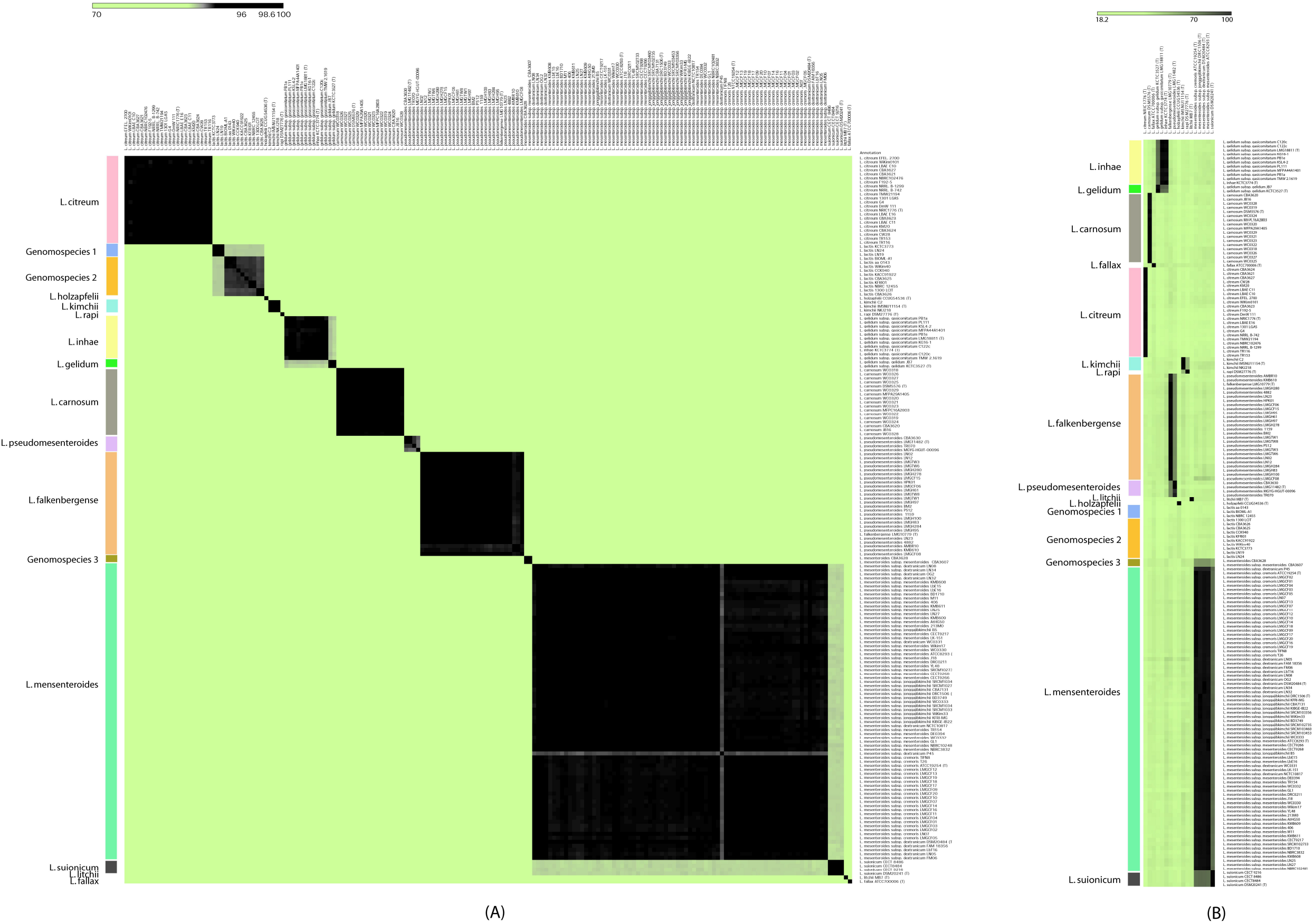
Whole genome-based similarity assessment using OrthoANI and dDDH. A). All is to all ANI similarity heatmap showing separate groups obtained by using a cut-off of 96%. B). Heatmap of digital DNA-DNA hybridization all across the type strains with the cut-off of 70% genome similarity.

**Figure 3:**
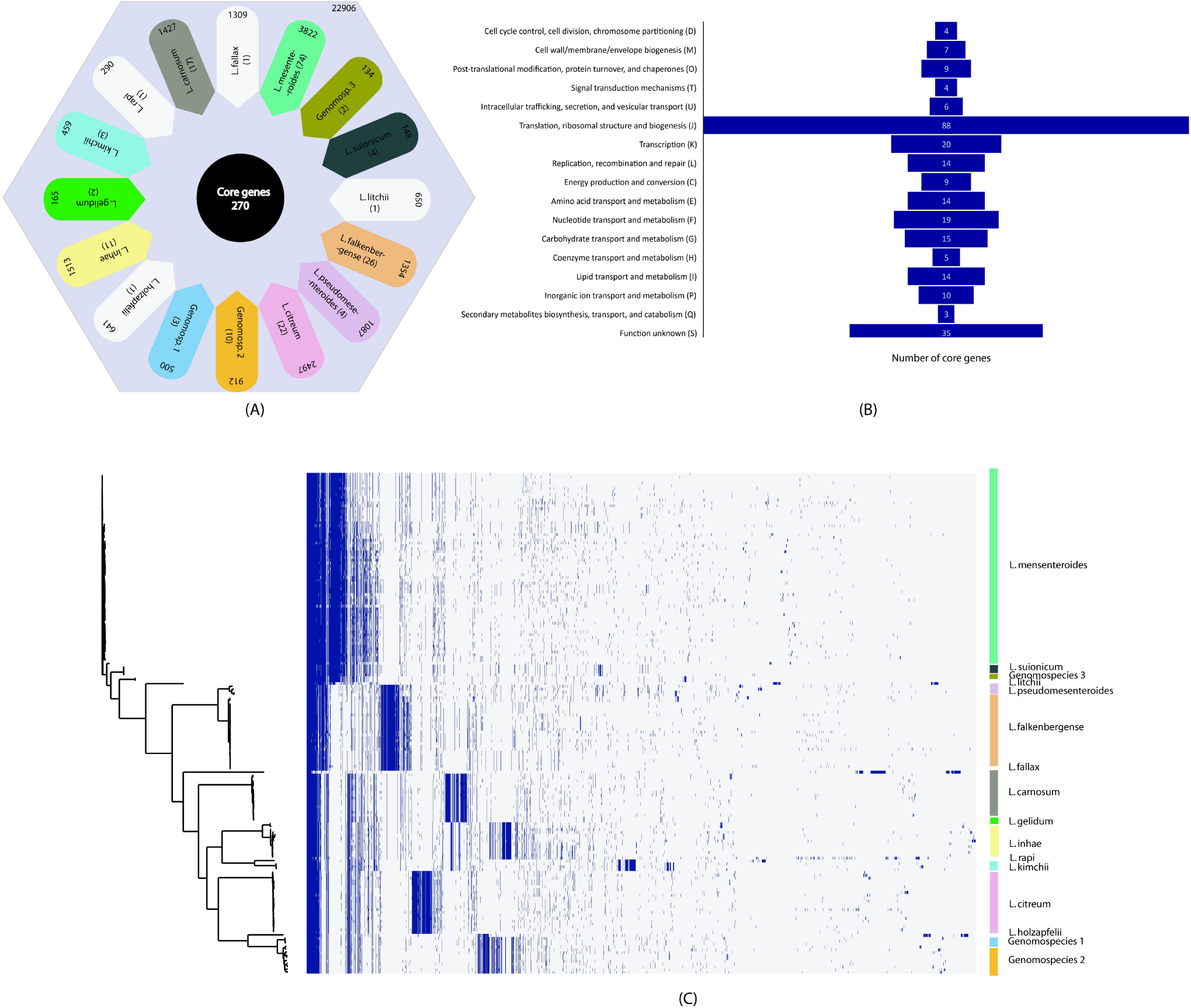
Pan-genome analysis: A). Pan-genome representation of all the gene clusters obtained from roary (22906). Core gene (270), and species-specific genes (unique gene) along with the number of strains in the group is marked in each group of genera *Leuconostoc*. B) Distribution of COG-based functional categories of core genes of the *Leuconostoc* genus. Here, the x-axis represents the number of genes and y-axis represents the functional categories. The count of gene cluster is indicated across classes in the box plot. C). Pan-genome plot representing the gene presence-absence across the strains. Core genome phylogroups are represented in the left isolates.

### Plasmid identification, antibiotic resistance and unique functional genomic attributes

To obtain the presence of plasmid in the strains of genus *Leuconostoc*, abricate v1.0.1 (https://github.com/tseemann/abricate) was implemented using plasmidFinder v2.1 database [55]. Antibiotic resistance genes were also checked using the card module v3.08 [56], ResFinder v4.1 [57] and ARG-ANNOT [58] of abricate. Genomic attributes for the prediction of biosynthetic gene clusters were identified using antiSMASH v6 [59]. All the type strains and the novel genomospecies were checked for the presence of a biosynthetic gene cluster.

### Carbohydrate active enzyme analysis

Carbohydrate-active enzymes (CAZy enzymes) (http://www.cazy.org/) among all the type strains and the novel genomospecies obtained from genome similarity analysis were identified using dbCAN2 [60]. dbCAN2 scans the genome using Hidden Markov model (HMM) profile which uses HMMdb v7 [61] (e-value of < 1e-15, coverage> 0.35), DIAMOND [62] (e-value < 1e-102) and Hotpep [63] (frequency > 2.6, hits > 6) to improve the prediction. The genes were annotated by at least two of the methods were taken for further analysis. The detailed carbohydrate-active enzyme family information was obtained on the CAZyme (http://www.cazy.org/). Identified CAZymes were broadly classified as glycoside hydrolase (GHs), glycosyltransferases (GTs), carbohydrate esterases (CEs), carbohydrate-binding enzymes (CBM), auxiliary active enzymes (AAs), and polysaccharide lyases (PLs).

## Result and Discussion

### Genomic features of the genus *Leuconostoc*

A total of 182 genomes of the genus *Leuconostoc* were used for the present study. Their average genome size ranged from 1 mb to 2.5 mb with an average GC content of 37%. The genomic attributes including #tRNA, #rRNA, #CDS etc. are summarized in table 1.

### Phylogenomics of the genus *Leuconostoc* reveals major phylogroups

Due to high similarity 16S rRNA based phylogeny could not resolve the constituent species of the genus *Leuconostoc*. For instance, *L. inhae* and *L. gelidum* subsp. *gasicomitatum; L. rapi* and *L. kimchii; L. falkenbergense* and few strains of *L. pseudomesenteroides; L. mesenteroides, L. suionicum, L. litchi* and *L. fallax* could not form distinct clade (Supplementary Figure 1). To get more accurate phylogeny, we obtained core genome-based phylogeny using roary and PhyloPhlAn (Figure 1, Supplementary Figure 2). Both of these approaches resulted in major reshuffling across the *Leuconostoc* sp. Strains of major species of genus *Leuconostoc* namely, *L. mesenteroides, L. pseudomesenteroides, L*. g*elidum* and *L. lactis* were not forming species-specific clades. Rather, each of these four species were splitting into two distinct phylogroups.

Out of 76 strains of *L. mesenteroides* 74 formed a distinct phylogroup along with the *L. mesenteroides* subsp. *mesenteroides* ATCC8293^T^ [1] confirming their species status. However, two of the remaining isolates namely, *L. mesenteroides* subsp. *mesenteroides* CBA3607 and *L. mesenteroides* CBA3628 formed a distinct phylogroup. Similarly, out of 29 strains of *L. pseudomesenteroides* 4 strains including *L. pseudomesenteroides* LMG11482^T^ [26] could be distinguished from the remaining 25 strains. Interestingly, these 25 strains of *L. pseudomesenteroides* formed a phylogroup with *L. falkenbergense* LMG10779^T^ [34]. Further, out of 12 strains *of L. gelidum* only one strain *L. gelidum* subsp. *gelidum* JB7 was forming phylogroup with *L. gelidum* subsp. *gelidum* KCTC3527^T^ [64]. Whereas, remaining 10 strains (*L. gelidum* subsp. *gasicomitatum*) were forming a phylogroup with the *L. inhae* KCTC3774^T^ [65]. Similarly, 13 strains of *L. lactis* were also forming two distinct phylogroups comprising of 10 and 3 strains respectively. However, due to the poor genome quality of the *L. lactis* type strain, we could not include in the present analysis. Overall, we could deduce 16 distinct phylogroups including 4 distinct unary strains which are representatives of previously described species namely, *L. fallax* ATCC700006^T^ [66], *L. rapi* DSM27776^T^ [27], *L. holzapfelii* CCUG54536^T^ [67], *L. litchii* MB7^T^ [67]

### Taxonogenomic assessment led to the identification of the novel genomospecies and reassignment of strains

Earlier taxonomic classification of species of *Leuconostoc* was mostly based on the classical taxonomy [1]. Description of the major species of genus *Leuconostoc* such as *L. mesenteroides* [1], *L. citreum* [26], *L. carnosum* [64], *L. inhae* [65] etc. was devoid of the whole genome-based approach such as ANI, dDDH and AAI etc., due to which, the current taxonomic classification of the *Leuconostoc* suggests them as a valid species (https://lpsn.dsmz.de/genus/leuconostoc). Robust phylogenomics of the genus *Leuconostoc* revealed several major reshufflings which need to be further confirmed by taxonogenomics.

Our whole genome-based taxonomic evidence using ANI (OrthoANI and fastANI) and dDDH correlated with the phylogenomics. For instance, three phylogroups consisting of earlier defined *L. lactis* and *L. mesenteroides* strains were found to be forming three novel genomospecies (GS1, GS2 and GS3) (Figure 2, Supplementary Figure 3). In addition to novel genomospecies, some of the previously defined strains of *L. pseudomesenteroides* and *L. gelidum* were identified as *L. falkenbergense* and *L. inhae* respectively. The remaining species stands valid in accordance with the standing in nomenclature (https://lpsn.dsmz.de/genus/leuconostoc) (LPSN link, table 2).

**Table: 2:**
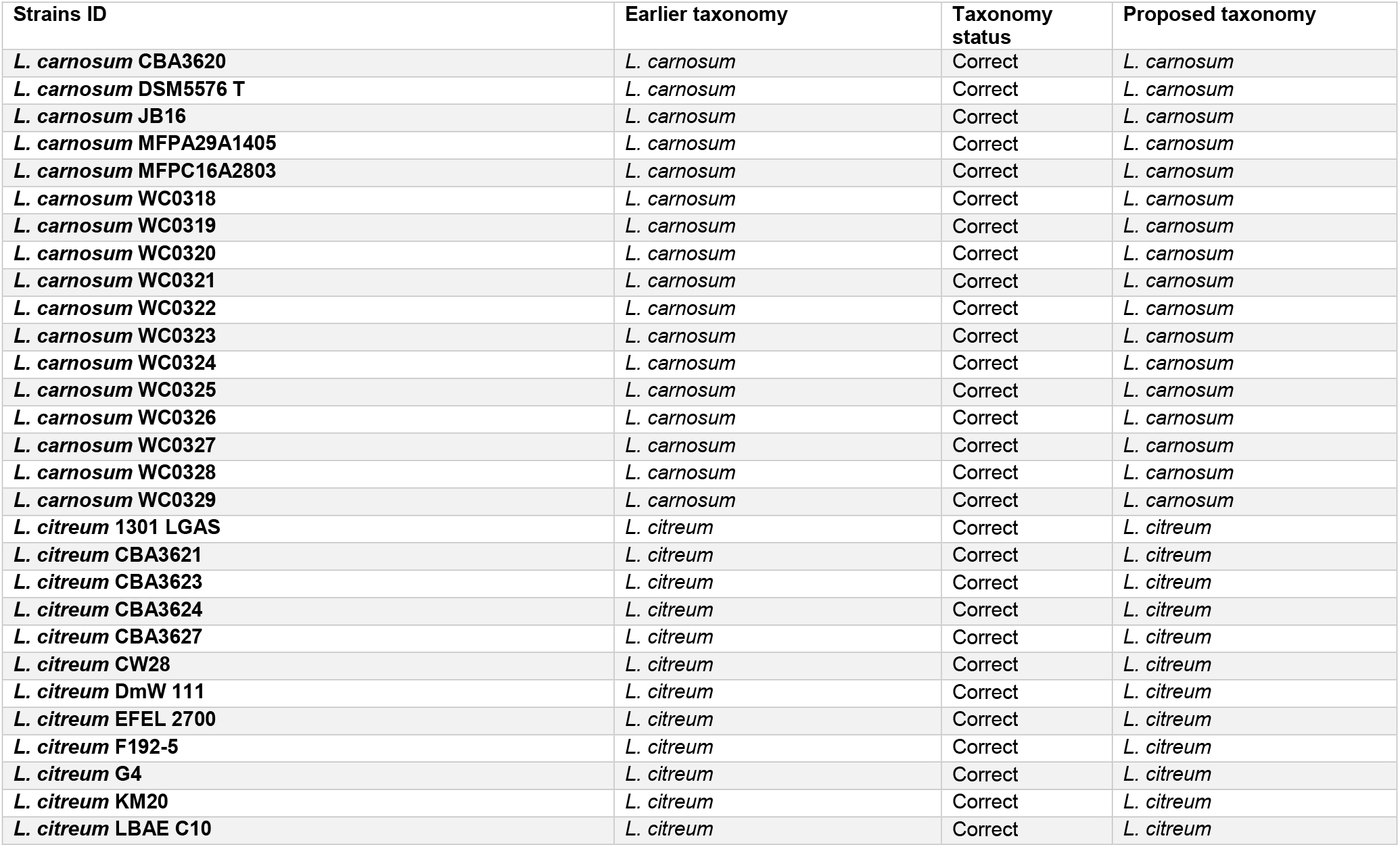
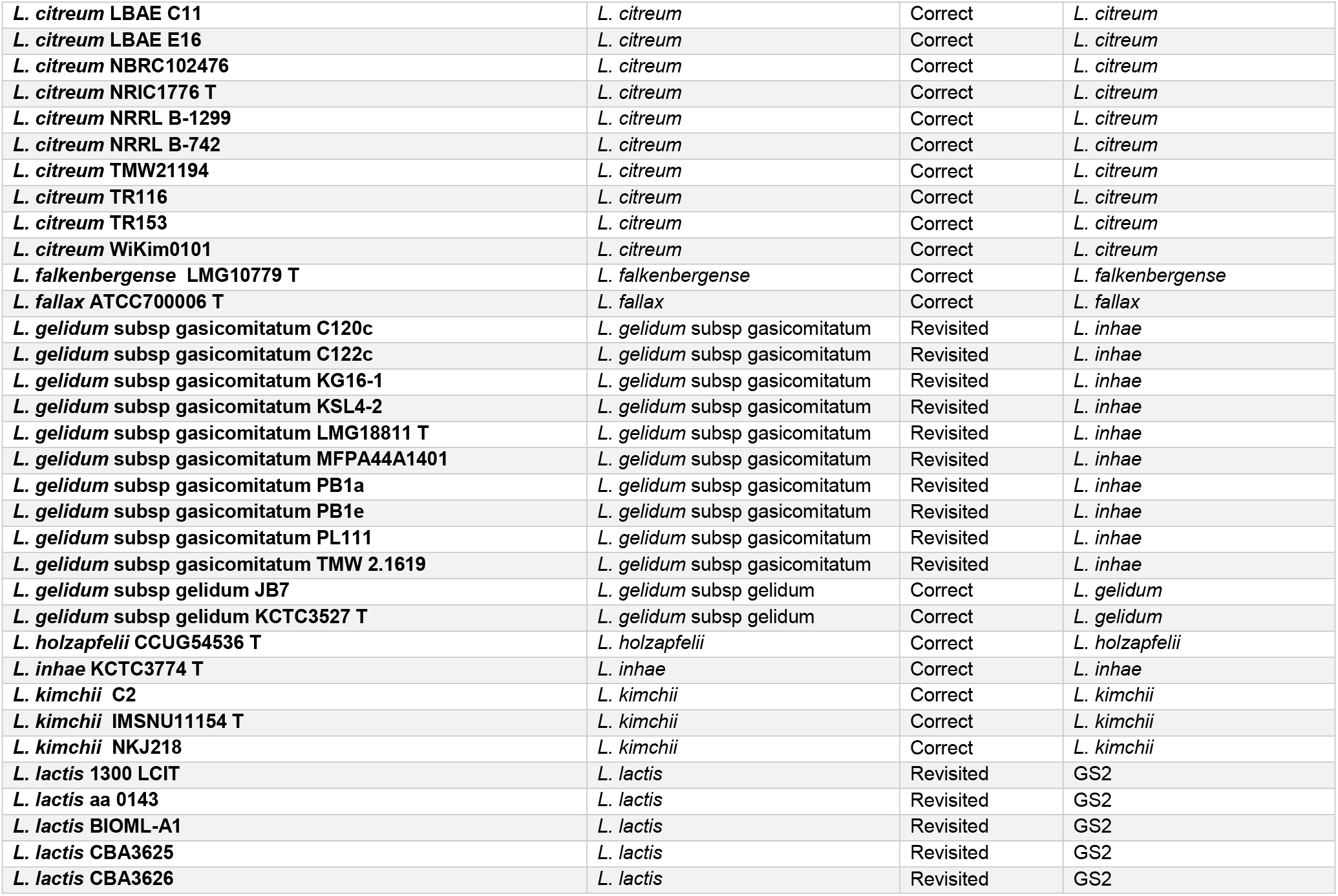
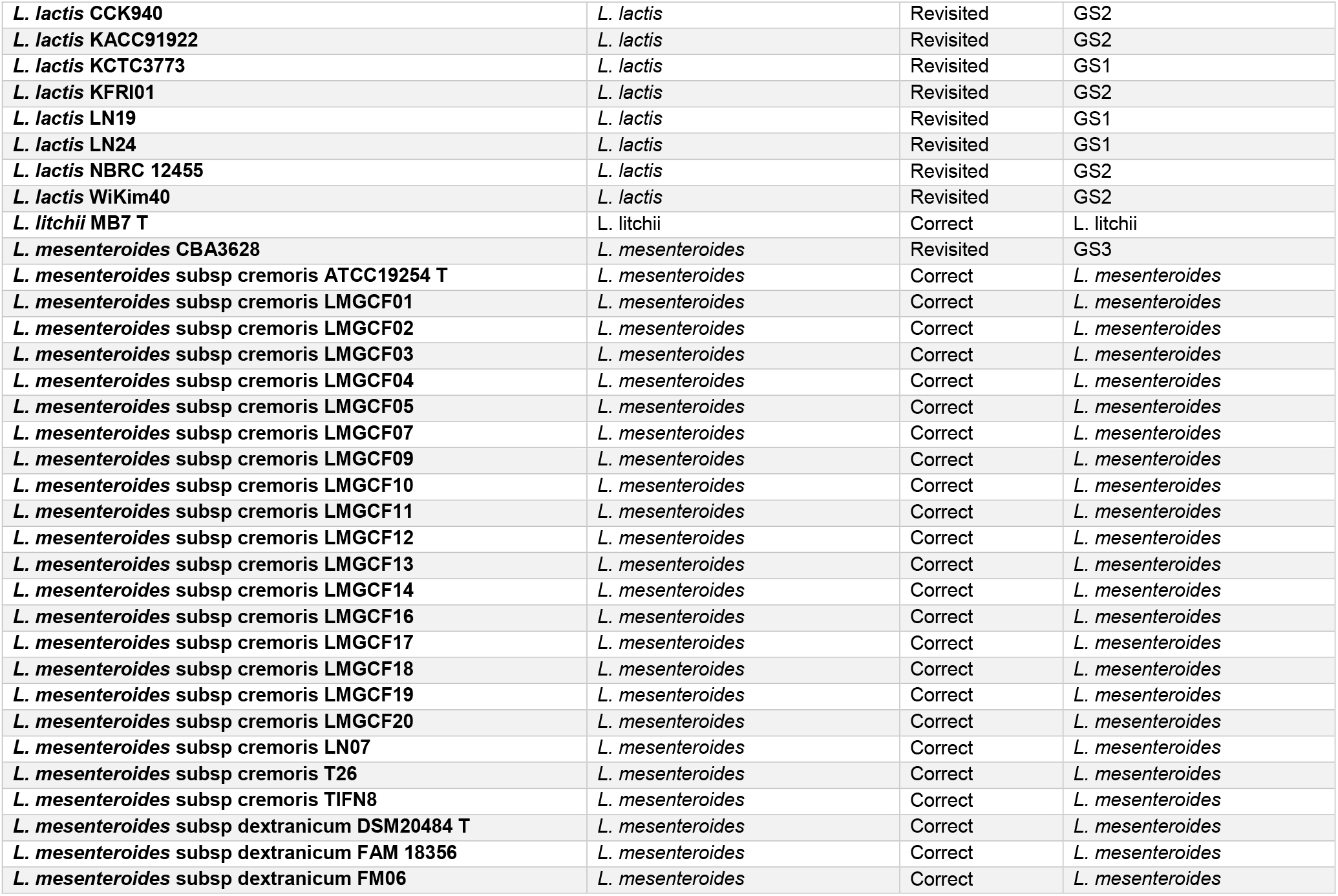
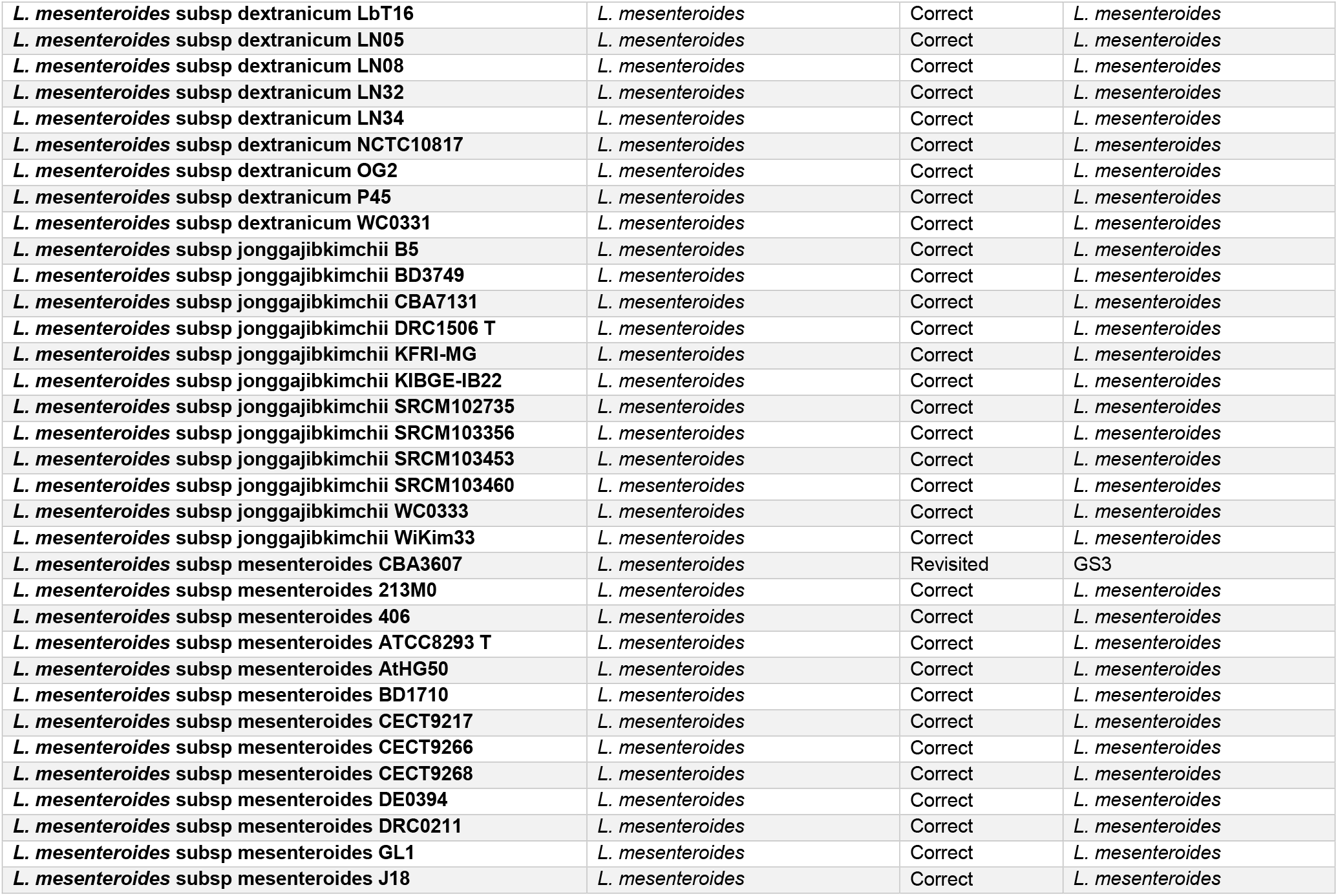
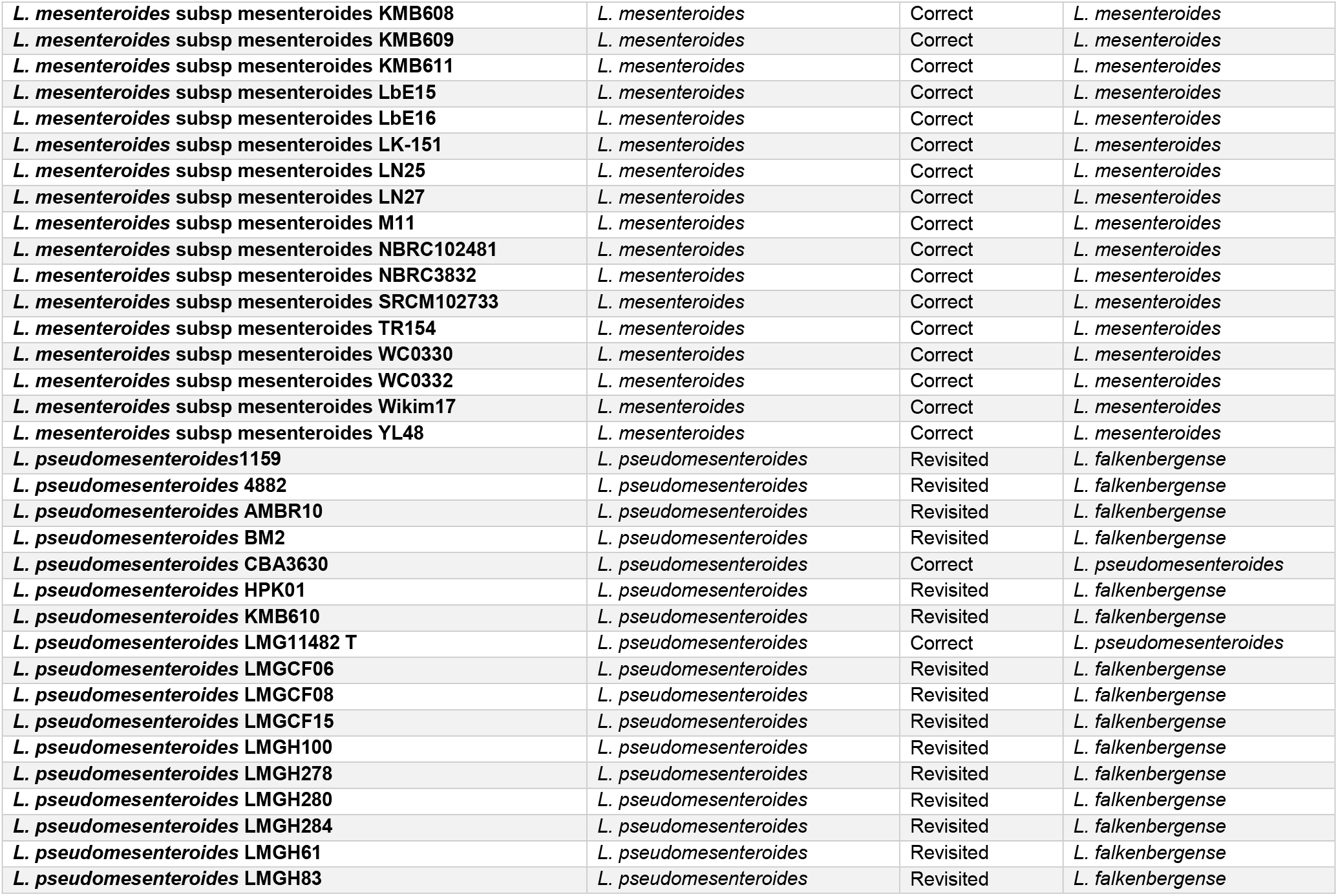
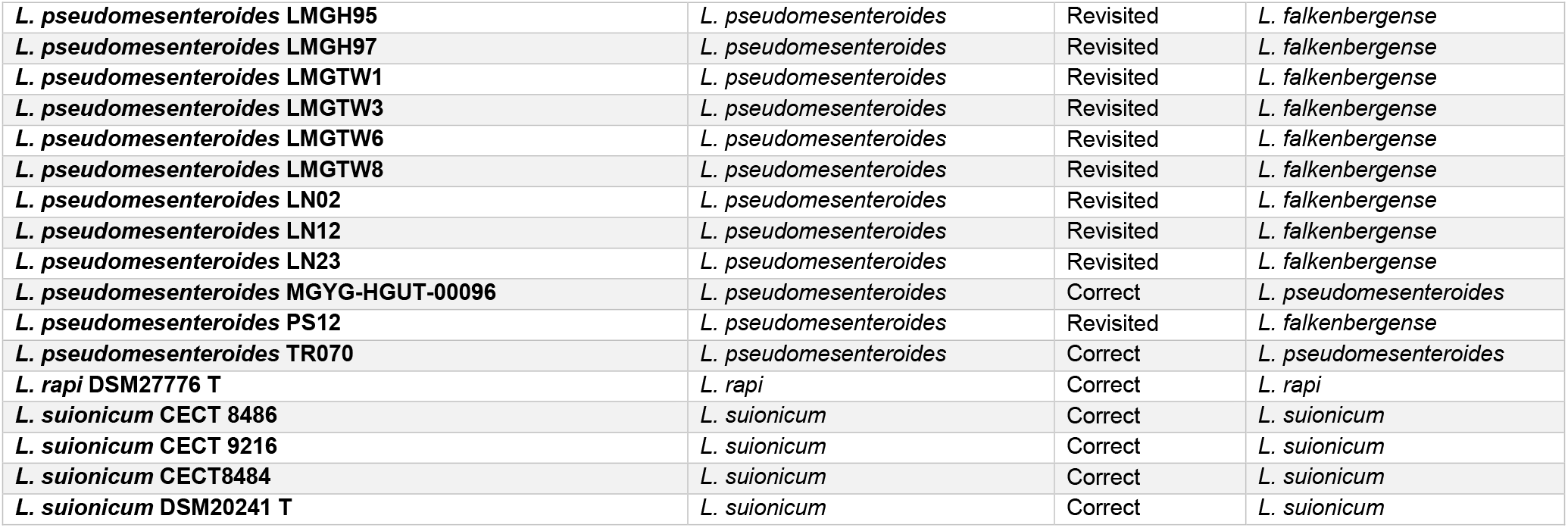
A descriptive list of the earlier classification and robust reclassification of the strain based on this study.

### Pan genome-based identification of core genome and species-specific genes

Pan-genome analysis using roary resulted in a comparatively small core genome (270 genes) and a large pan genome (22906 genes) which reveals a wide diversity of gene content within genus *Leuconostoc*. Here, we could get unique genes specific to the 16 species identified using taxonogenomics and phylogenomics (Figure 3A). *L. mesenteroides* (3822 genes) and *L. citreum* (2497 genes) largely contributed to the diversity of the pangenome. While, GS3 (134 genes), *L. suionicum* (148 genes) and *L. gelidum* (165 genes) had the least species-specific genes. Interestingly, amongst the unary species members, *L. fallax* showed the highest diversity with 1309 genes. The gene clusters obtained suggested a high number of accessory genes (shell and cloud) of 22,559 depicting a high degree of diversity across the species of genus *Leuconostoc*. Pan-genome matrix as well depicted the presence of species-specific genes (Figure 3C). Functional classification of the core genes suggested the majority of the information storage and processing (44%) and metabolism (32%) related genes (Figure 3B).

### Unique genomic attributes across spp. of *Leuconostoc*

High diversity in the genus *Leuconostoc* is evident from the gene content analysis. On the contrary, they were found to harbour pLCK4 (12183 bp) and LkipL4719 (21924 bp) plasmids in 97 and 86 genomes respectively. Plasmid pLCK4 was originally derived from *L. citreum* KM20 [68], which is one among the four high copy number plasmids of *L. citreum* KM 20 [68]. Plasmid pLCK4 which is considered to be a part of pMBLT00 [69], is considered to be a shuttle vector for inducing overproduction of D-lactate in *Leuconostoc* and *Lactococcus lactis* strains [69]. LkipL4719 is first reported in the *Leuconostoc kimchii* IMSNU 11154 [70] which harbours several metal transport gene families, suggesting its intrinsic nature across the species of *Leuconostoc* [70]. Additionally, 8 different plasmids namely pKLC4 (36602) in 11 genomes, pIGMS32 (or ColRNAI_1) (9294 bp) in 6 genome, pJB01 (2235 bp) in genome and pLFE1 (4031 bp) in 2 genomes. Whereas plasmid pYSI8 (4973 bp), pMG828-1 (Col(MG828) _1) (1902 bp), pJE1 (5149 bp) and pBI143 (2747 bp) are present in one genome each (Figure 1). In addition to low diversity in the plasmid, antibiotic resistance genes were also present but only in 4 genomes out of 182 *Leuconostoc* strains analysed using CARD, resfinder and arg-annot database integrated in abricate (supplementary table 2). However, genomic analysis of the *Leuconostoc* human pathogens is limited due to a lack of genomic resources.

Whereas, we found Type III polyketide synthase (PKSs) in all the type strains of 16 species identified (Figure 4 A). Type III polyketide synthase (PKSs) are homodimers of ketosynthase which catalyze the condensation of one or several molecules of extender substrate onto a starter substrate through an iterative decarboylative Claisen condensation reactions [71, 72]. Microbial type III PKSs seem to use an acyl–acyl carrier protein (ACP) as a starter substrate, [73], and in some cases, type III PKS genes form a cluster with ACP or fatty acid biosynthetic genes [74, 75]. Future studies on type III PKSs will provide important insights into the properties of these enzymes and their role in the biosynthesis of natural products. Moreover, strains of genus *Leuconostoc* is reported to harbour several bacteriocins belonging to class IIa bacteriocins, leucocin H, mesentericin etc. [76–79] imparting antimicrobial activity against foodborne pathogens [80]. In-depth genomic investigation of bacteriocins and antimicrobial peptides are the need of the hour to explore the potential of the species of genus *Leuconostoc*.

**Figure 4:**
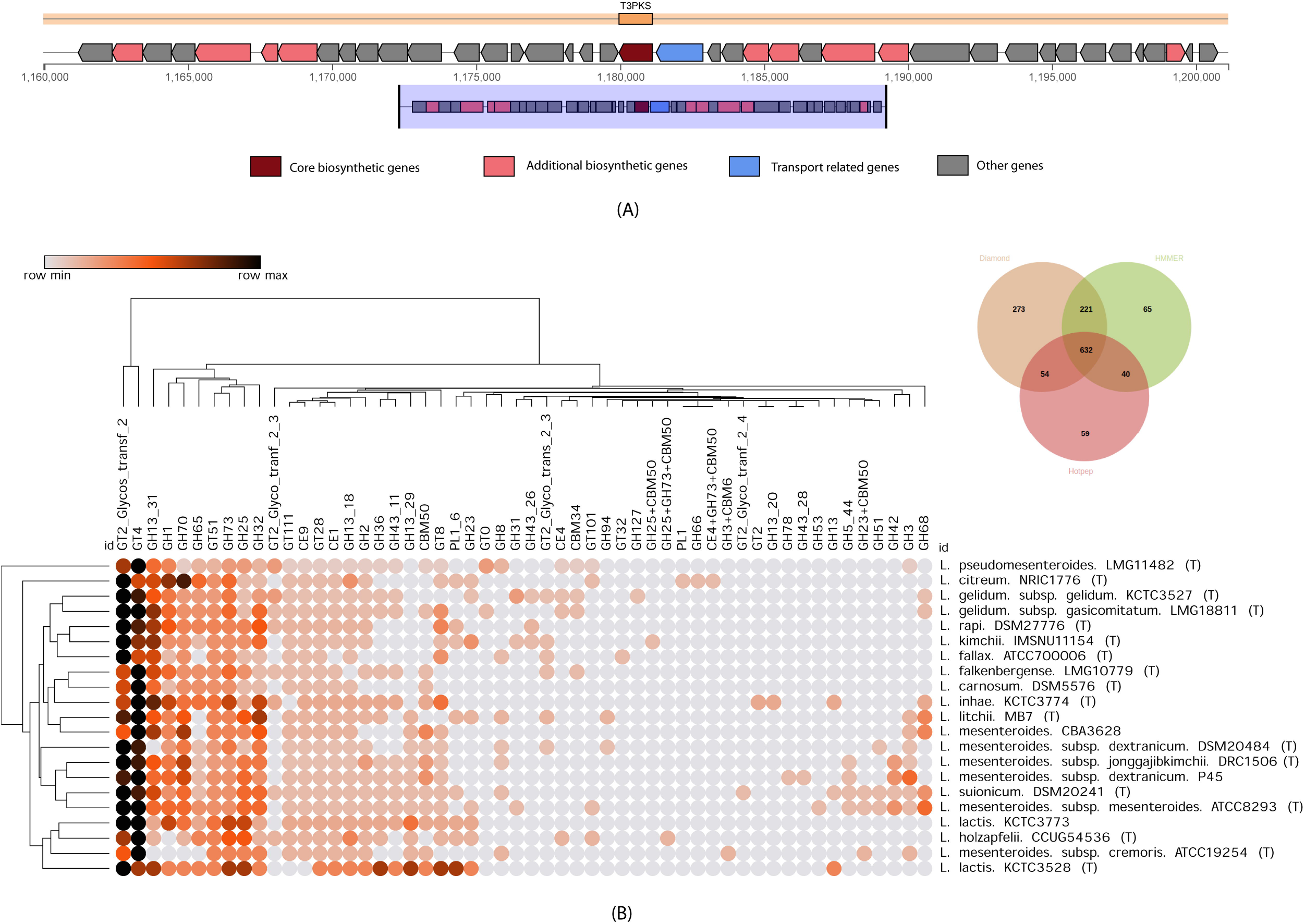
Type III PKS and CAZymes: A). The presence of type III PKS gene cluster was present in all the type strains in the study. The presence of the core biosynthetic gene in almost all the type strains was identified. Other genes such as additional biosynthetic genes, transporter genes etc. were also identified. B). Heatmap showing all the different classes of the carbohydrate-active enzyme classified under GH: Glycoside Hydrolase; CE: Carbohydrate Esterase; PL: Polysaccharide Lyase; GT: Glycosyltransferase; AA: Auxiliary Activities; CBM: Carbohydrate-Binding Module. Venn diagram revealing carbohydrate-active enzyme identified by three different approaches of HMMER, diamond and hotpep. Heatmap was generated based on the presence of the carbohydrate-active enzyme by at least two approaches.

### Carbohydrate-Active Enzyme (CAZymes)

Among type strains of 16 phylogroups identified using genome similarity and phylogenomic assessment reveals the presence of CAZyme groups of GH, GT, CR, AA, CBM and PL using dbCAB2. These CAZymes groups are the enzymes involved in the degradation, modification, and creation of glycosidic bonds. Altogether 1344 protein-coding genes were detected by one of three methods namely hotpep, Diamond and HMMER (Figure 4 B). We found all the major CAZyme groups distributed across the strains represented with a heatmap (Figure 4 B) reveals the differential presence of all the classes of CAZyme in the strains of *Leuconostoc*. Most of the identified gene clusters were distributed across GH (52%), GT (40%), CE (4.75), CMB (2.2) and PL (1%) families. (Summarized in supplementary table 1). We found diverse GHs namely GH13 (7.07%), GH70 (5.49%), GH73 (5.38%), GH1 (5.12%), GH 32 (4.85%), GH25 (4.54%), GH65 (2.95%) across *Leuconostoc* sp. GHs catalyse the cleavage of O-glycosidic bonds linking the carbohydrate moieties or carbohydrate and non-carbohydrate moieties. GH13 family of CAZymes reveals the presence of amylases, beta-glucosidases, beta-xylosidases, beta-galactosidase gene cluster across the species of *Leuconostoc*. Similarly, the GH70 family includes transglucosylases which catalyse the intra- or intermolecular replacement of glycoside molecules at the anomeric position, giving rise to new glycoside molecules, such as oligosaccharides [81]. GH73 family includes N-acetylmuramidase which cleaves the β-1,4-glycosidic bond between the N-acetylglucosaminyl (GlcNAc) and N-acetylmuramyl (MurNAc) moieties in bacterial peptidoglycan [82]. Glycosyl transferases are primarily involved in the formation of glycosidic bonds by transfer of sugar moiety from the activated sugar donor to the acceptor molecule. We identified three major GTs namely GT4 (13.5%), GT2 (12.6%) and GT51 (4.3%). GT4 a glycosyltransferase family 4 protein is responsible for accessory Sec system glycosyltransferase GtfA. GT2 glycosyltransferase family 2 protein is responsible for undecaprenyl-diphospho-muramoyl pentapeptide and beta-N-acetylglucosaminyltransferase and putative glycosyltransferase, exosortase G system-associated. GT51 is responsible for PBP1A family penicillin-binding protein and murein polymerases. These are majorly involved in the synthesis of the peptidoglycan cell wall and play crucial roles in maintaining the integrity of the cell wall [83]. Likewise, CAZymes prevent under CE, CBM and PL categories are CE9 (2.11%), CE1 (2.1%), CBM50 (1.79%) and PL1 (1%).

## Conclusion

Several taxonomic reclassifications were found by the present study inferring the true phylogenetic positioning of the strains of genus *Leuconostoc* (Table 2). True phylogeny will help future research to better identify and reclassify the strains of the genus belonging to biotechnologically important genera. In-depth genomic investigations of the strains suggest the presence of several determinants such as plasmid and type III PKS system etc. and the absence of antibiotic gene cluster also approves GRAS status. However, minor reports of *Leuconostoc* strains as human pathogens have raised concerns about their biotechnological implications. The present phylo-taxonogenomic study provides a robust taxonomic framework of the genus which will be valuable in the identification of these clinically relevant strains. Such future genomic investigations of strains from diverse niches including nosocomial, environment, food and industry will shed light on their niche-specific attributes. This will be critical in surveillance to demarcate safe to use strains for food and industry from their pathogenic counterparts. Thus, our work is expected to promote research on the biotechnological important genus which is long overlooked and to better understand the intrinsic property of these important microbes.

---

**Supplementary Figure 1: 16S rRNA tree:** A phylogenetic tree based on ML-based obtained by16S rRNA gene sequences of all the strains under study. Bootstrap values are represented by blue dots. Strain name of some of the complete genome which resulted in multiple copies of 16S rRNA were appended with 1, 2, 3 etc.

**Supplementary Figure 2: Phylogenetic tree obtained using PhyloPhlAn:** A phylogenetic tree based on more than 400 conserved genes. Bootstrap values are marked with blue dots.

**Supplementary Figure 3: fastANI:** Heatmap obtained using genome similarity matrix obtained using fastANI.

**Supplementary Table 1:** A descriptive list of antibiotic resistance genes predicted using several databases such as using CARD, resfinder and arg-annot.

**Supplementary Table 2:** A descriptive list of CAZyme identified using dbCAN2.

## Abbreviations

GH: Glycoside Hydrolase
CE: Carbohydrate Esterase
PL: Polysaccharide Lyase
GT: Glycosyltransferase
AA: Auxiliary Activities
CBM: Carbohydrate-Binding Module
ANI: Average Nucleotide Identity
dDDH: digital DNA-DNA hybridization
LAB: Lactic Acid Bacteria
Type III PKS: Type III polyketide synthase

