## Supplementary figures and images for "Phylo-taxonogenomics of 182 strains of genus *Leuconostoc* elucidates its robust taxonomy and biotechnological importance"

### Supplementary Figure 1

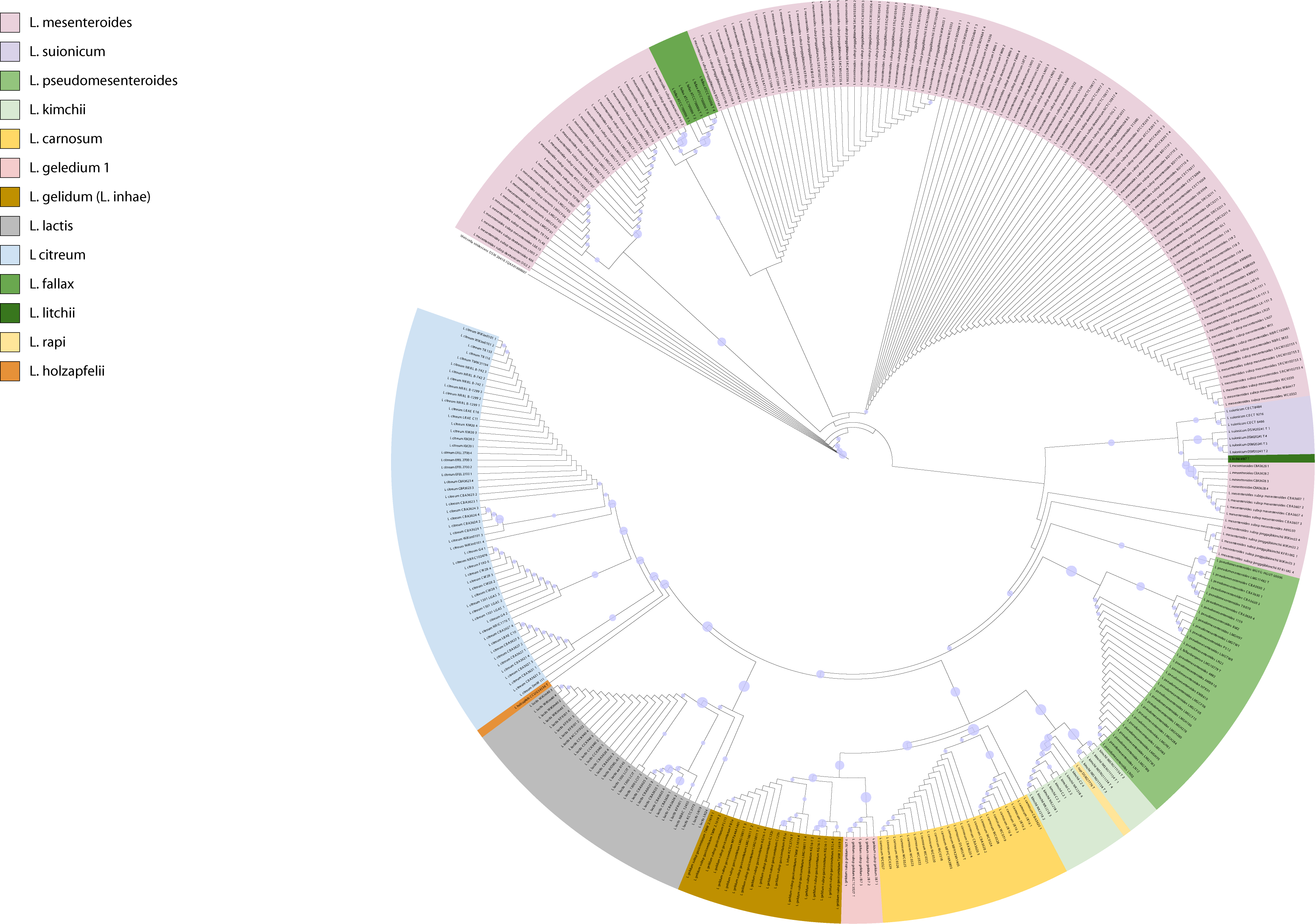

### Supplementary Figure 2

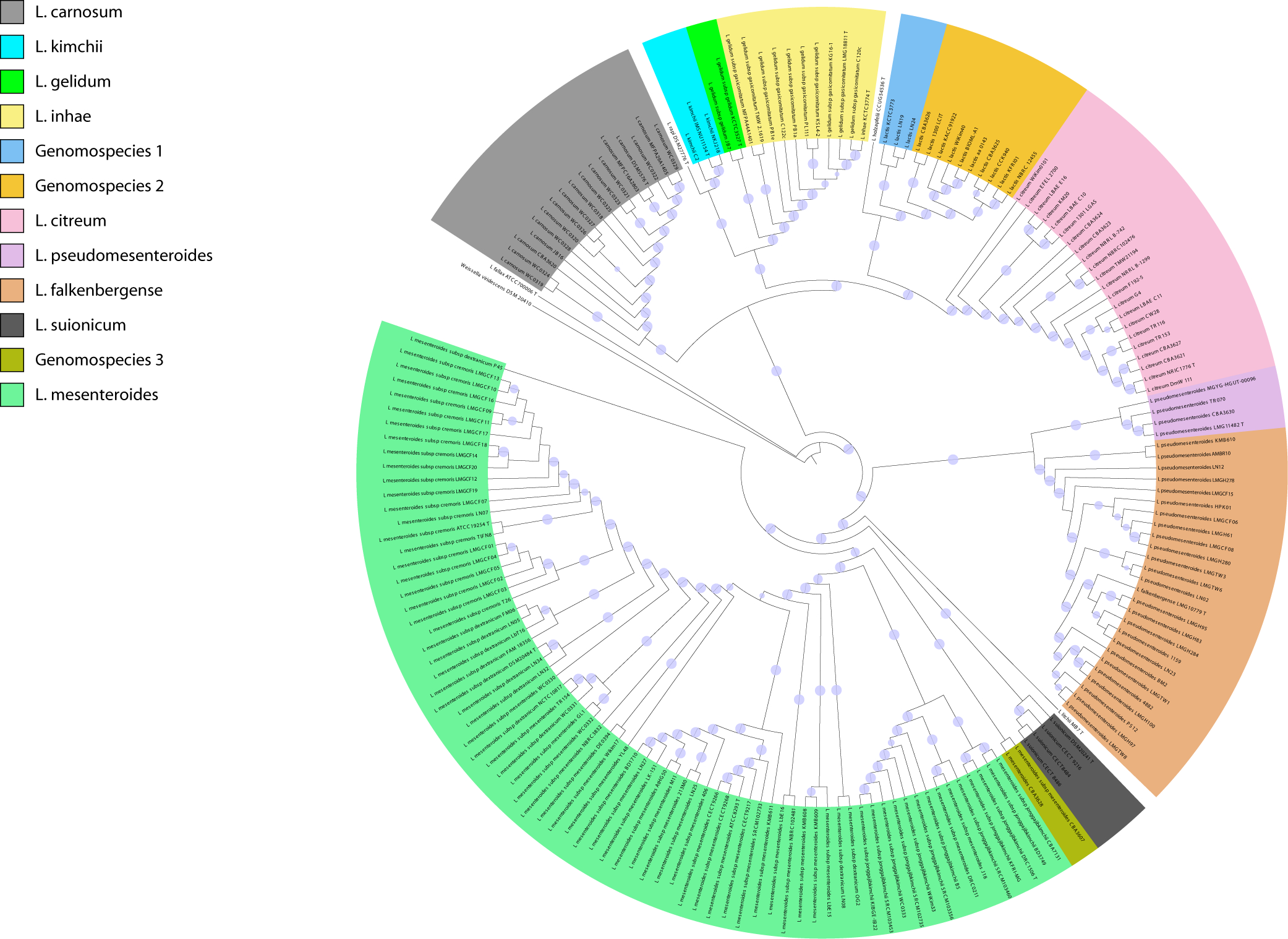

### Supplementary Figure 3

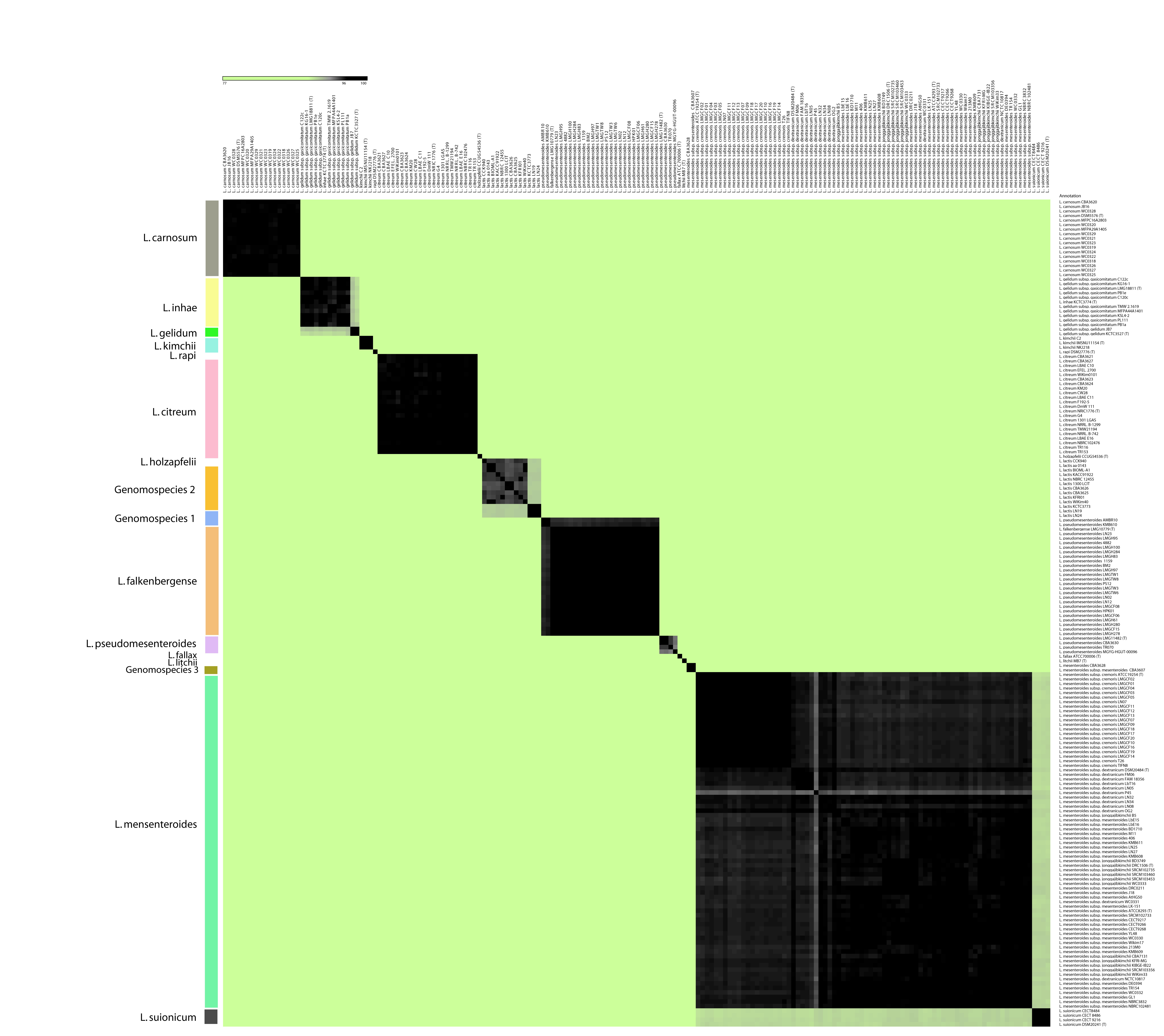
